# RCOR1 promotes myoblast differentiation and muscle regeneration

**DOI:** 10.1101/2024.05.27.596049

**Authors:** Martina Pauk, Fan Wang, Petri Rummukainen, Mauricio Ramm, Hanna Taipaleenmäki, Riku Kiviranta

**Affiliations:** Institute of Biomedicine, University of Turku, Turku, Finland; Institute of Musculoskeletal Medicine, University Hospital, LMU Munich, Munich, Germany; Musculoskeletal University Center Munich, University Hospital, LMU Munich, Munich; Department of Molecular Medicine and Surgery, Karolinska Institutet, Stockholm, Sweden; Department of Endocrinology, Division of Medicine, University of Turku and Turku University Hospital, Turku, Finland

**Keywords:** RCOR, Lysine specific demethylase 1 (LSD1), Myoblast, Skeletal muscle regeneration, Myogenesis

## Abstract

RCOR proteins belong to a family of highly conserved transcription corepressors (RCOR1, RCOR2 and RCOR3) that regulate the activity of associated histone demethylase 1 (LSD1) and histone deacetylase 1/2 (HDAC 1/2) in chromatin-modifying complexes. Despite the described function of LSD1 in skeletal muscle differentiation and regeneration, the role of RCOR family in myogenesis remains unknown. We found that RCOR1 is highly expressed in proliferating myoblasts and activated satellite cells, but not in mature myofibers during postnatal growth and regeneration of skeletal muscle. Knockdown of RCOR1 impaired myoblast differentiation and fusion by inhibiting the expression of the key myogenic regulatory factor myogenin. Moreover, RCOR1 depletion impaired myoblast proliferation through increasing the expression of cell cycle inhibitor p21. Consistently, in a mouse model of skeletal muscle injury, depletion of RCOR1 supressed satellite cell activation and differentiation which resulted in impaired muscle regeneration. RCOR1 was found physically associated with LSD1 and myogenic regulatory factor MyoD and contributed to LSD1 stability in myoblasts. As for other RCOR family members, RCOR2 had no effect on myoblast differentiation while the loss of RCOR3 increased myoblast proliferation leading to supressed expression of myogenic markers MyoD and myogenin and impaired myoblast differentiation. However, germline deletion of RCOR3 (RCOR3^-/-^) did not affect muscle phenotype, suggesting a possible functional redundancy among RCOR family members during muscle development. Together, our findings indicate that RCOR1 acts in concert with LSD1 as a novel positive regulator of myogenesis and skeletal muscle regeneration.

## Introduction

Skeletal muscle is the most abundant and highly complex tissue of the vertebrate body. The process of muscle formation known as myogenesis, occurs during embryonic development, postnatal growth and regeneration of adult skeletal muscle upon injury. Skeletal muscle has remarkable regenerative capacity mainly due to its resident population of Pax7-expressing muscle stem cells called satellite cells (SCs)^1^. During adult myogenesis, SCs are activated and form mononucleated myoblasts that exit the cell cycle, subsequently fusing into multinucleated myotubes and ultimately form mature muscle fibers^1^. These processes are regulated by a complex molecular network including several transcription factors, among which four myogenic regulatory factors (MRFs), MyoD, Myf5, Mrf4 and myogenin, have a vital role in controlling muscle-specific gene expression^1,2^. MyoD, Myf5 and Mrf4 act as the early muscle determination factors^3–5^, while myogenin, induced by the other three MRFs, regulates the later stages of myogenic differentiation and the myofiber maturation^6,7^. MRFs also cooperate with other transcriptional regulators such as myocyte enhancer factor 2 (MEF2) to regulate and activate muscle specific target genes which are required for sarcomeric organization and contractility such as myosin heavy chain (MyHC)^8^.

The epigenetic mechanisms involved in the regulation of myogenesis are not fully understood. Several studies have demonstrated an essential role of lysine-specific histone demethylase 1 (LSD1) in skeletal muscle differentiation and regeneration. LSD1 acts as a corepressor by demethylating mono- and di-methylated lysine 4 and/or 9 residues of histone H3 (H3K4me1/2 and H3K9me1/2), leading to either transcriptional repression or activation, respectively^9–11^. Inhibition of LSD1 has been shown to impair skeletal myoblast differentiation in C2C12 cells^12–14^. A proposed mechanism of action suggests direct binding of LSD1 and MyoD to the myogenin promoter, leading to demethylation of di-methylated form of H3K9 and consequent activation of myogenic genes^12^. In addition, LSD1 was shown to be necessary for the timely expression of MyoD through the regulation of the MyoD core enhancer during muscle differentiation^15^. Finally, LSD1 was identified as a key epigenetic regulator of muscle regeneration by repressing brown adipocyte differentiation of SCs and promoting myogenesis^16^.

LSD1 is found in association with the transcriptional corepressor CoREST complex, whose primary components are LSD1, histone deacetylase 1 or 2 (HDAC1/2) and RCOR1, 2 or 3 scaffold protein^17^. Within this complex, the presence of RCOR protein is essential for the nucleosome recognition and histone tail binding leading to effective demethylation and deacetylation by LSD1 and HDAC1/2, respectively^9,17^. There are three vertebrate RCOR proteins encoded by separate genes: RCOR1 (CoREST1), RCOR2 (CoREST2) and RCOR3 (CoREST3). Although all three RCORs share high sequence similarity and ability to interact with LSD1, they possess differential biochemical properties and distinct molecular actions. RCOR2 and RCOR3 display lower transcriptional repressive capacity than RCOR1, which could be due to the variations in their structures and altered protein-protein interactions^18^. Accordingly, RCOR1 and to a lesser extent RCOR2 efficiently stimulate LSD1-mediated demethylation of nucleosomes, while this was not observed for the LSD1-RCOR3 complex in erythroid cells^9,19–21^. This led us to hypothesise that RCOR1 may be the primary RCOR protein that acts in concert with LSD1 in myoblasts and is involved in regulation of the myogenic gene program. RCOR1 has been studied in a wide variety of cell types, and shown to play important roles in mouse erythropoiesis^22^, neuronal gene regulation^23,24^ and tumorigenesis^25^ by modifying chromatin structure. However, its potential role in myogenesis has not yet been investigated.

Here, we demonstrated that RCOR1 expression is linked to the progression of myogenic differentiation, postnatal growth and muscle regeneration upon injury. Furthermore, RCOR1 knockdown significantly impaired myoblast proliferation and differentiation by supressing the expression of cell cycle regulators and key myogenic factors, respectively. RCOR1 depletion in muscle reduced activation and differentiation of SCs, leading to an impaired muscle regeneration following injury. By investigating other members of the RCOR family, we found that RCOR2 had no effect on myoblast differentiation, while absence of RCOR3 inhibited the expression of key myogenic factors and the differentiation of myoblasts. However, RCOR3 was dispensable for muscle development, suggesting a possible functional redundancy among other members of the RCOR family during muscle development. Collectively, we have identified for the first time the specific role of RCOR1 in skeletal muscle differentiation and regeneration.

## Results

### RCOR1 is dynamically expressed during myoblast differentiation and *in vivo* myogenesis

To determine whether RCOR1 is regulated during myogenesis, we analysed its expression and localization patterns in several myogenesis systems *in vitro* and *in vivo*. First, we used the mouse skeletal muscle cell line C2C12, which represent a well-established model of the skeletal muscle differentiation process. Differentiation was initiated upon serum deprivation leading C2C12 myoblasts to fuse into multinucleated myotubes during a 5-day period. RCOR1 mRNA expression remained unchanged during this period (Fig 1A). However, immunoblotting showed that RCOR1 protein gradually declined throughout myoblast differentiation, along with an increase in the abundance of myogenic markers Myogenin and MyHC (Fig. 1B). Interestingly, LSD1 mRNA expression moderately increased during differentiation, but protein abundance gradually decreased following the onset of myogenic differentiation (Fig. 1A and B), suggesting that both LSD1 and RCOR1 are downregulated during myogenesis through yet unknown mechanisms. To confirm these findings in primary cells, we isolated primary myoblasts from 8±1 week-old male mice and induced them to differentiate to myotubes over a 3-day period. Consistently, both RCOR1 and LSD1 mRNA and protein decreased during the differentiation of primary myoblasts (Fig. 1C and D). In order to analyse the cellular distribution of RCOR1, we performed immunofluorescence staining in C2C12 myoblasts and primary myoblasts. RCOR1 was mainly expressed in the nuclei of myoblasts and differentiated myotubes in both cell culture models (Fig. 1E and S1A), which is in accordance with its role in chromatin modification. Nuclear localization of LSD1 in C2C12 cells was also confirmed by immunostaining (Fig. S1B). In addition, we investigated the expression levels of other members of RCOR family, RCOR2 and RCOR3, during differentiation by qPCR. Similar to RCOR1, RCOR2 and RCOR3 mRNA expression remained unchanged during the C2C12 myoblast differentiation but decreased following the onset of primary myoblast differentiation (Fig 1F and G). Interestingly, RCOR1 was expressed significantly higher than RCOR3 in C2C12 myoblasts and showed highest abundance in primary myoblasts compared to RCOR2 and RCOR3 (Fig. S1C).

**Figure 1.**
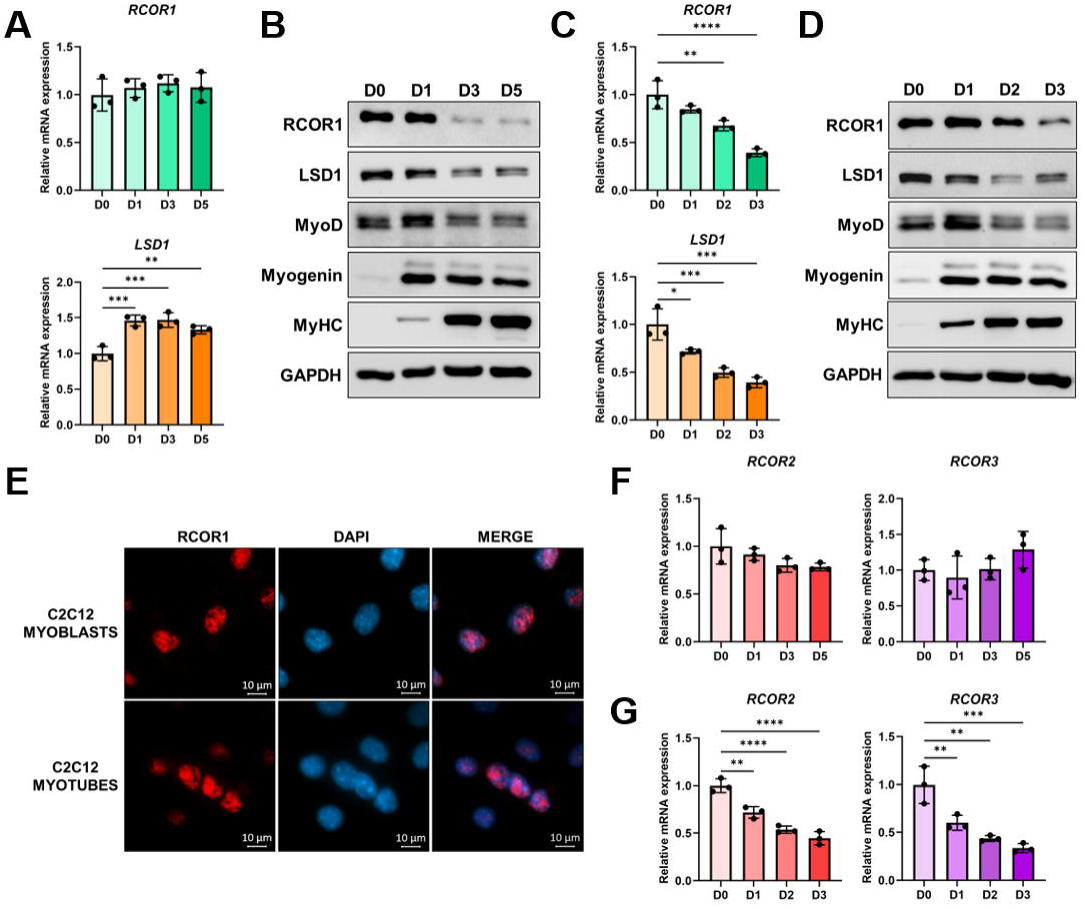
RCOR1 is dynamically expressed during myoblast differentiation. C2C12 myoblasts cultured in growth medium (GM) were induced to differentiate by switching to differentiation medium (DM) for 5 days. The expression levels of RCOR1, LSD1, myogenic regulatory factors (MRFs) and MyHC were detected in C2C12 cells by (**A**) qPCR and (**B**) immunoblotting and in primary mouse myoblasts by (**C**) qPCR and (**D**) immunoblotting. (**E**) Immunofluorescence staining for RCOR1 was performed in C2C12 myoblasts and myotubes (red, RCOR1; blue, DAPI). (**F**) The expression levels of RCOR2 and RCOR3 were detected in (**F**) C2C12 cells and (**G**) primary myoblasts by qPCR. Pictures are one representative of 3 independent experiments. Data are presented as means ± SD; n=3 mice per group, * P < 0.05; ** P < 0.01; *** P < 0.001, **** P < 0.0001.

Next, we investigated the expression of RCOR1 during myogenesis *in vivo*. Both RCOR1 and LSD1 were highly expressed in limb muscles of newborn mice on postnatal day 1 (P1) but decreased in 2- and 4-week-old mice (Fig. 2A and B). Consistently, key myogenic markers MyoD and myogenin showed a similar expression pattern during muscle development with lower abundance in adult muscles, as expected (Fig. 2A and B). To explore the expression of RCOR1 during skeletal muscle regeneration, TA muscles were injured by CTX injection and allowed to regenerate for 10 days. Four days post injury, we observed extensive inflammatory cell infiltration accompanied by small regenerating myofibers with centrally located nuclei (Fig. S2A). After 10 days of regeneration, the CTX-injured muscle appeared nearly fully regenerated with only small number of interstitial cells left. Both RCOR1 and LSD1 mRNA and protein expression were highly increased in CTX-treated TA muscle relative to contralateral PBS-treated muscle at 4 days following injection (Fig. 2C and D). At 7-10 days postinjury, RCOR1 and LSD1 expression gradually declined but remained detectable over the course of regeneration. RCOR1 was observed in mononucleated cells at the injury site and within newly formed muscle fibers in CTX-injured muscle (Fig. 2E). Importantly, myogenic markers MyoD and myogenin showed similar dynamics in the expression, consistent with their role in coordinating myogenesis (Fig. 2C and D). Consistent with previous findings in myoblasts, we found that RCOR1 is the most abundant RCOR family member in CTX-injured muscle (Fig. S2B). Collectively, these data revealed that RCOR1 is associated with active myogenesis *in vitro* and *in vivo* and might play a role in skeletal muscle development and regeneration.

**Figure 2.**
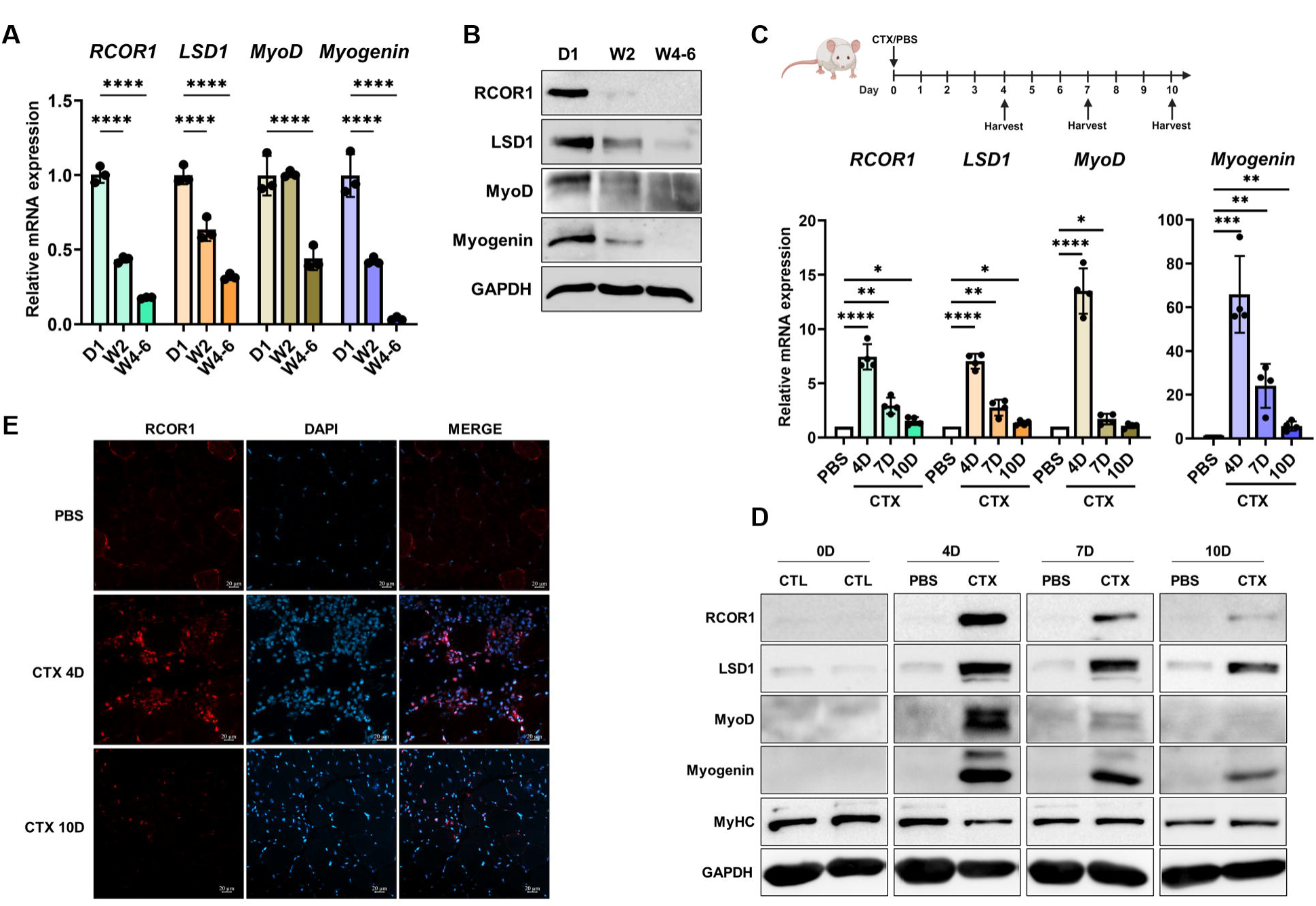
RCOR1 is dynamically expressed during *in vivo* myogenesis. Hindlimb muscles from mice at postnatal day 1 (D1), week 2 (W2) and week 4-6 (W4-6) were analysed by (**A**) qPCR and (**B**) immunoblotting to detect mRNA and protein levels of RCOR1, LSD1 and MRFs. TA muscles were injured with Cardiotoxin (CTX) or PBS injection and harvested on day 4, 7 and 10 for (**C**) qPCR and (**D**) immunoblotting analysis of RCOR1, LSD1, MRFs and MyHC expression. (**E**) Upon cryosection, immunofluorescence staining for RCOR1 (red) and DAPI (blue) in TA muscles was performed at day 4 and 10 following CTX injury. Pictures are one representative of 3 independent experiments. Data are presented as means ± SD; n=3 mice per group, * P < 0.05; ** P < 0.01; *** P < 0.001, **** P < 0.0001.

### RCOR1 regulates myoblast differentiation

To investigate whether RCOR1 is required for skeletal myogenesis, we knocked-down RCOR1 expression in C2C12 myoblasts by using small interfering RNAs (siRNAs). Knockdown efficiency was verified by qPCR and immunoblotting (Fig. S3A). As shown by MyHC immunostaining, RCOR1 deficiency severely impaired myogenic differentiation and fusion as indicated by a lower differentiation and fusion index relative to control siRNA (Fig. 3A). siRCOR1-transfected myoblasts remained as individual mononucleated cells and did not fuse to form multinucleated myotubes. Consistently, the expression of myogenic regulatory factor myogenin and muscle structural protein MyHC was significantly reduced at early and late stages of differentiation, respectively (Fig. 3B and C). Immunofluorescence staining of myogenin revealed a reduced number of myogenin positive myoblasts following RCOR1 knockdown, supporting the inhibitory effect on the differentiation (Fig. S3B). Of note, the expression of myogenic regulatory factor MyoD remained unchanged during differentiation of C2C12 myoblasts (Fig. 3B and C). In primary cultures, siRCOR1 transfected myoblasts exhibited smaller and fewer myotubes relative to elongated multinucleated myotubes formed by control myoblasts (Fig. S3C). Consistent with the observed decrease in myotube formation, fusion index of siRCOR1 transfected primary myoblasts was significantly reduced as well as mRNA levels of myogenin, Mrf4 and MyHC compared to control (Fig. S3C and D). However, no difference was observed in the differentiation index and at the protein level of myogenic regulatory factors and MyHC (Fig. S3C and D). Together, these data indicate that RCOR1 is essential for myoblast differentiation and fusion in C2C12 myoblasts.

**Figure 3.**
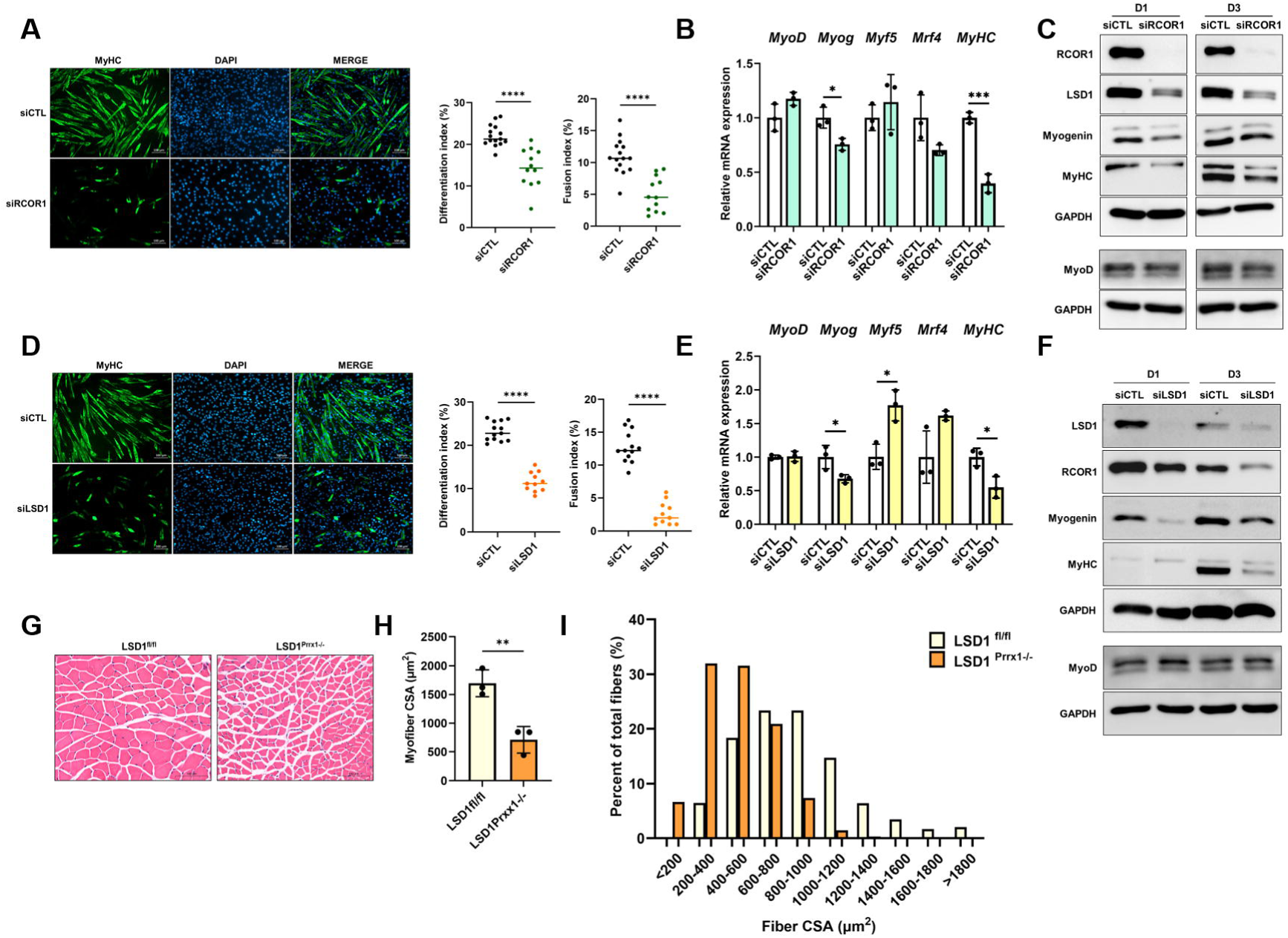
RCOR1 regulates myoblast differentiation. C2C12 cells transfected with control siRNA (siCTL), RCOR1 or LSD1 siRNA (siRCOR1; siLSD1) were cultured in growth medium (GM) and induced to differentiate by switching to differentiation medium (DM) for 3 days. (**A**) Terminally differentiated myotubes transfected with siRCOR1 were visualized by anti-MyHC immunofluorescent staining (green, MyHC; blue, DAPI) at day 3 in DM. The differentiation index and fusion index were counted. (B) qPCR and (**C**) immunoblotting were performed to detect mRNA and protein levels of RCOR1, LSD1 and myogenic regulatory factors (MRFs) at day 1 and MyHC at day 3 in DM. (**D**) Terminally differentiated myotubes transfected with siLSD1 were visualized by anti-MyHC immunofluorescent staining (green, MyHC; blue, DAPI) at day 3 in DM. The differentiation index and fusion index were counted. (**E**) qPCR and (**F**) immunoblotting were performed to detect mRNA and protein levels of RCOR1, LSD1 and myogenic regulatory factors (MRFs) at day 1 and MyHC at day 3 in DM. TA muscles of control (LSD1fl/fl) and LSD1 knockout mice (LSD1Prrx1-/-) (4 weeks of age) were stained by (**G**) H&E and analysed for (**H**) mean myofiber cross-sectional area (CSA) and (**I**) percent distributions of fiber cross-section area CSA (μm2). Data are presented as means ± SD; n=3 mice per group. * P < 0.05; **, P < 0.01; ***, P < 0.001, **** P < 0.0001.

As described above, RCOR1 is a part of the CoREST complex containing LSD1 and HDAC1/2^9,17,26^. Moreover, RCOR1 directly binds to LSD1 and regulates its activity and function^27,28^. Previous studies indicated that LSD1 plays a role in myogenic differentiation^12–14^. Similarly, we observed that LSD1 knockdown significantly impaired myoblast differentiation and fusion as shown by MyHC immunostaining in C2C12 cells and primary myoblasts (Fig. 3D and S4A). Furthermore, the loss of LSD1 reduced the expression of myogenin and MyHC (Fig. 3E and F and Fig. S4B). However, contrary to previous reports^12–14^, LSD1 had no impact on MyoD mRNA and protein expression (Fig. 3E and F and Fig. S4B). Different functional approaches for LSD1 depletion and inactivation in mice have demonstrated the requirement of LSD1 in myogenic differentiation and muscle regeneration^16,29,30^. Here, we crossed LSD1^fl/fl^ mice with Prrx1-Cre mice to specifically delete the LSD1 gene in limb bud mesenchymal progenitors (LSD1^Prrx1-/-^)^31^. Due to the severe limb phenotype and lower body weight^31^, LSD1^Prrx1-/-^ mice were analysed at 4 weeks of age to minimize the suffering of the animal. Likewise, H&E-stained TA muscles revealed significant decrease of myofiber cross-sectional area (CSA) and altered fiber size distribution in LSD1^Prrx1-/-^ mice (Fig. 3G-I). CSA measurement revealed a higher proportion of small diameter fibers in LSD1^Prrx1-/-^ mice, while larger fibers were less represented relative to control (Fig. 3I), suggesting that targeted deletion of LSD1 in the limb bud progenitors resulted in defective muscle development and postnatal growth.

RCOR1 belongs to RCOR family of proteins consisting of two more members. Both RCOR2 and RCOR3 also associate with LSD1 and functional redundancy has been previously reported among the family members^18,21,32^. To determine whether other RCORs compensate for the loss of RCOR1, we first examined the expression of RCOR2 and RCOR3 following RCOR1 knockdown and found no significant changes in mRNA expression in C2C12 myoblasts (Fig. S5A). Next, we examined a potential role for RCOR2 and RCOR3 in the myogenic differentiation using siRNA-mediated knockdown (Fig. S5B). RCOR2 knockdown had no effect on myogenic differentiation and expression of key myogenic markers and muscle structural protein MyHC relative to control (Fig. 4A and B). Conversely, the loss of RCOR3 effectively blocked the formation of myotubes, as assessed by reduced MyHC immunofluorescence (Fig. 4C), decreased differentiation and fusion index (Fig. 4C) and downregulated expression of MyoD, myogenin and MyHC (Fig. 4D and E). The role of RCOR3 during myogenic differentiation was further confirmed in primary mouse myoblasts (Fig. S5C and D). To determine the function of RCOR3 in muscle development *in vivo*, we generated mice with a germ-line deletion of RCOR3 (*RCOR3^-/-^*) (Fig. S6). We observed no difference in body weight and *RCOR3^-/-^* mice showed normal muscle structure without any signs of degeneration or change in myofiber CSA distribution relative to *RCOR3^+/+^* animals (Fig 4F-H). These data indicate that RCOR3 is not required for muscle development *in vivo*.

**Figure 4.**
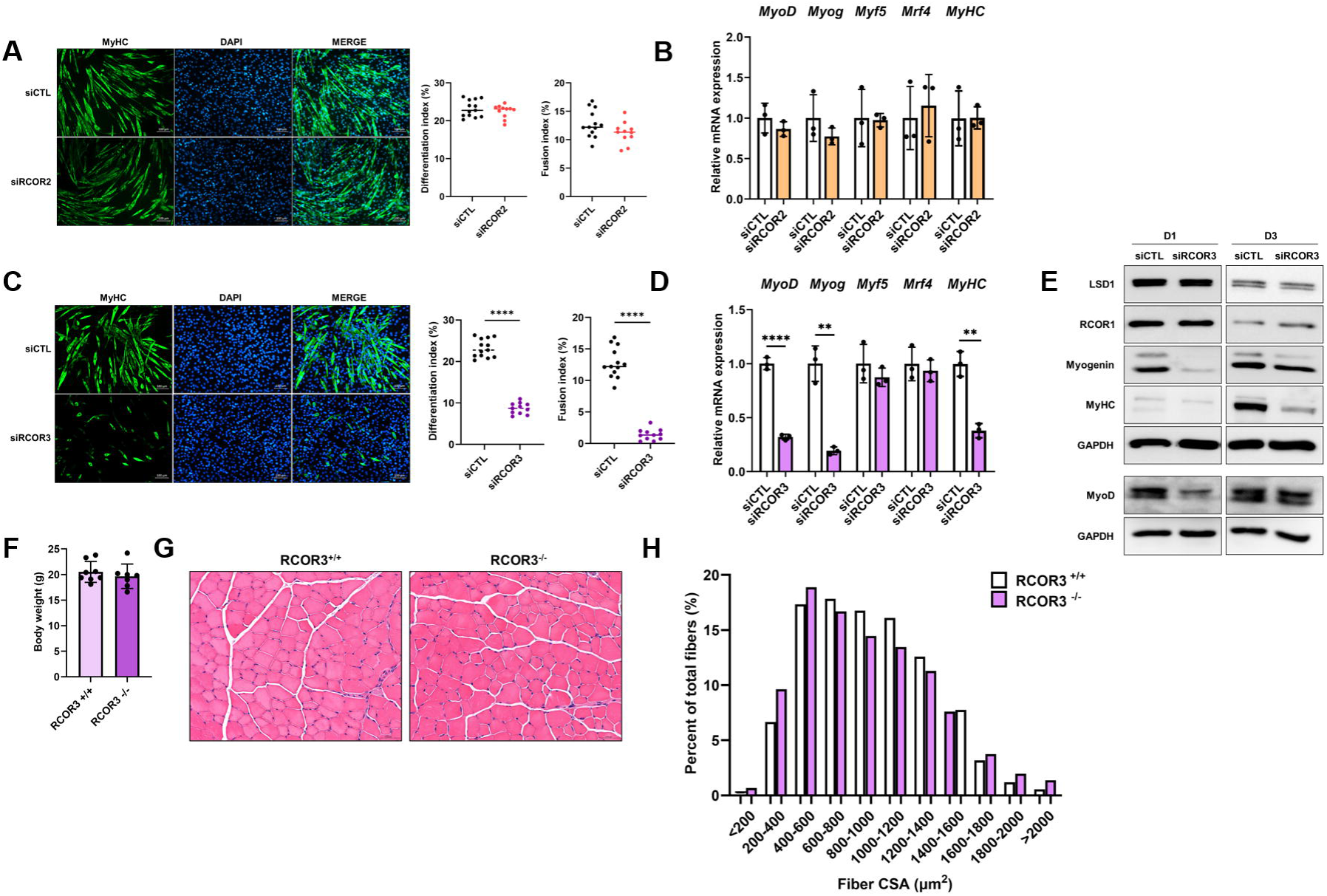
Redundancy between RCOR proteins in skeletal muscle differentiation. C2C12 cells transfected with control siRNA (siCTL), RCOR2 or RCOR3 siRNA (siRCOR2; siRCOR3) were cultured in growth medium (GM) and induced to differentiate by switching to differentiation medium (DM) for 3 days. (**A**) Terminally differentiated myotubes transfected with siRCOR2 were visualized by anti-MyHC immunofluorescent staining (green, MyHC; blue, DAPI) at day 3 in DM. The differentiation index and fusion index were counted. (**B**) qPCR was performed to detect mRNA levels of myogenic regulatory factors (MRFs) at day 1 and MyHC at day 3 in DM. (**C**) Terminally differentiated C2C12 myotubes transfected with siRCOR3 were visualized by anti-MyHC immunofluorescent staining (green, MyHC; blue, DAPI) at day 3 in DM. The differentiation index and fusion index were counted. (**D**) qPCR and (**E**) immunoblot were performed to detect mRNA and protein levels of RCOR1, LSD1 and myogenic regulatory factors (MRFs) at day 1 and MyHC at day 3 in DM. (**F**) RCOR3^+/+^ and RCOR3^-/-^ mice were analysed for the body weight at 6 weeks of age (n=5). TA muscles were stained by (**G**) H&E and analysed for (**H**) percent distributions of fiber cross-section area (CSA) (μm^2^). Pictures are one representative of 3 independent experiments. Data are presented as means ± SD. n=5 mice per group. * P < 0.05; **, P < 0.01; ***, P < 0.001, **** P < 0.0001.

### RCOR1 regulates myoblast proliferation

Differentiation and proliferation of skeletal muscle cells are mutually exclusive and highly regulated processes. Since cell cycle arrest is critical for muscle differentiation, we examined the effect of RCOR1 on C2C12 myoblast proliferation. RCOR1 knockdown reduced the number of cells cultured in both, growth and differentiation medium (Fig. 5A). Consistently, the metabolic rate, as measured by the MTT assay, decreased in siRCOR1 transfected myoblasts relative to controls (Fig. 5B). Next, we performed BrdU incorporation assay in C2C12 myoblasts and primary myoblasts and observed a decreased percentage of BrdU-positive nuclei upon RCOR1 knockdown in both, growth and differentiation medium relative to control cells (Fig. 5C and S7A), indicating a decrease in proliferation upon RCOR1 knockdown.

**Figure 5.**
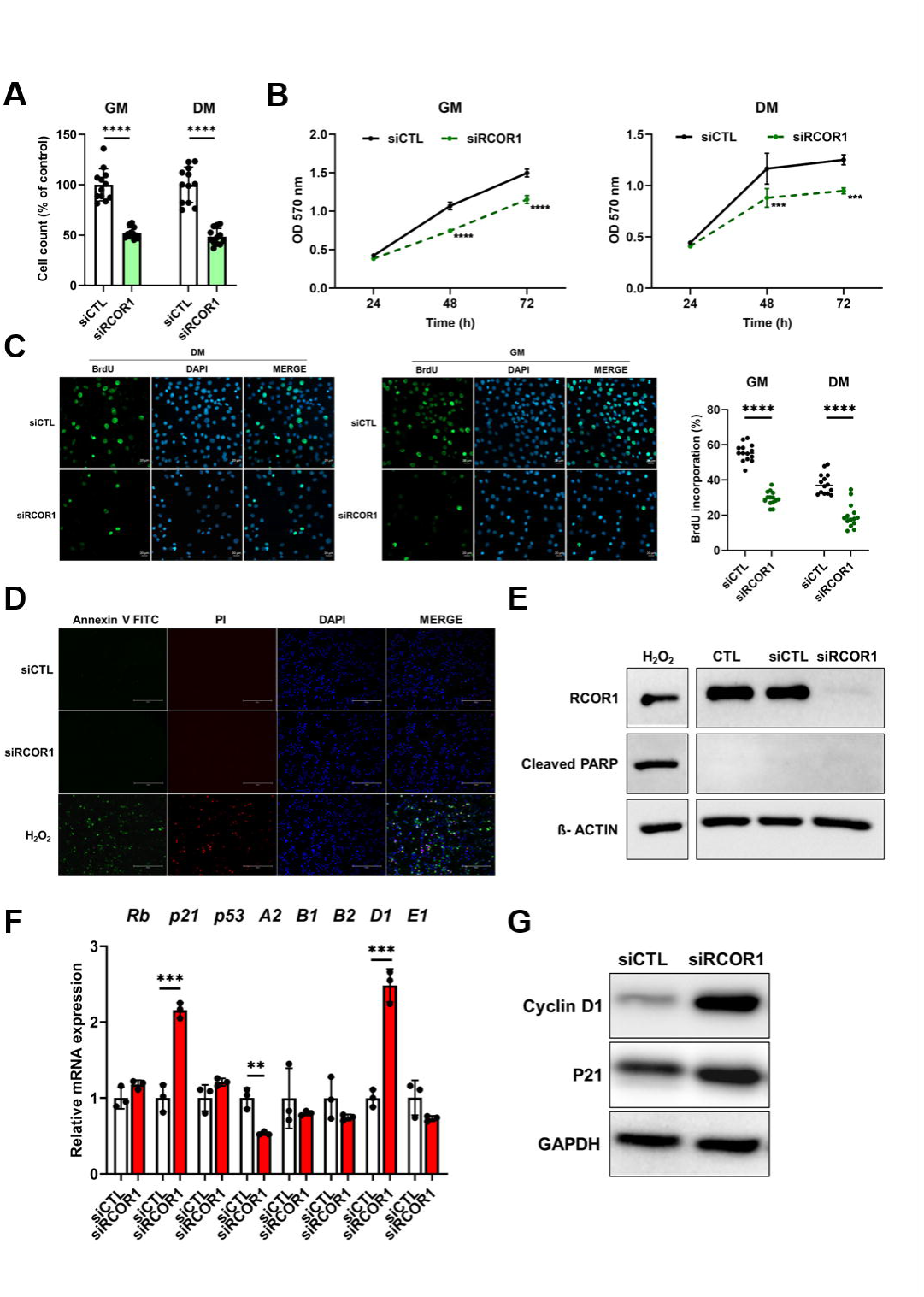
RCOR1 regulates myoblast proliferation. C2C12 cells transfected with control siRNA (siCTL) or RCOR1 siRNA (siRCOR1) were cultured in growth medium (GM) and then switched into fresh GM or differentiation medium (DM) for additional 1 day. (**A**) DAPI-stained nuclei were counted in GM and DM. (**B**) Proliferation analysis was assessed by MTT assay at indicated time points in GM and DM. (**C**) After 1 day in GM or DM, cells were stimulated with GM or DM containing 10 µM BrdU for 4 hours and then immunostained for the detection of BrdU incorporation (green BrdU; blue DAPI). The percentage of the number of BrdU positive cells was calculated. (**D**) Cell death of transfected C2C12 cells cultured in DM for 1 day was quantified by Annexin V and propidium iodide (PI) staining and (**E**) immunoblotting of cleaved PARP protein levels. For positive control, cells were treated with pro-apoptotic agent H_2_O_2_ (1 mM). Transfected C2C12 cells cultured in DM for 1 day were analysed for mRNA and protein levels of cell cycle regulators (Rb, p21, P53, Cyclin A2, Cyclin B1, Cyclin B2, Cyclin D1 and Cyclin E1) by (**F**) qPCR and (**G**) immunoblotting. Pictures are one representative of 3 independent experiments. Data are presented as means ± SD. * P < 0.05; **, P < 0.01; ***, P < 0.001, **** P < 0.0001.

To determine whether the decrease in myoblast proliferation was due to an increased apoptosis or altered cell cycle regulation, we performed Annexin-V-FITC/PI and DAPI staining. Immunofluorescence analysis revealed no difference between siCTL and siRCOR1-transfected cells and neither group showed significant apoptotic cell death (Fig. 5D). This result was further confirmed by immunoblotting of cleaved PARP, another marker of apoptosis (Fig. 5E). C2C12 cells treated with hydrogen peroxide (H2O2), a well-known apoptosis inducer, were used as a positive control and showed increased portion of annexin V-FITC positive/PI-positive cells and increased PARP cleavage (Fig. 5E).

Cell cycle progression is controlled by the activity of cyclins and their associated cyclin-dependent kinases (CDKs) whose activities are negatively regulated by CDK inhibitors (CKIs)^33^. We analysed the expression of several cell cycle regulators (cyclin A2, cyclin B1, cyclin B2, cyclin D1, cyclin E, Rb, p21 and p53) by qPCR in C2C12 myoblasts. siRCOR1-transfected myoblasts exhibited a reduction of cyclin A2, while cyclin D1 increased (Fig. 5F and G). The expression of p21 was also investigated. Consistent with reduced proliferation, RCOR1 knockdown significantly increased the mRNA and protein abundance of p21, which is a negative regulator of the cell cycle (Fig. 5F and G). Interestingly, LSD1 knockdown failed to affect C2C12 myoblast proliferation as determined by BrdU incorporation assay and protein expression of p21 (Fig. S7B and C), indicating that LSD1 is not essential for myoblast proliferation. Based on these results, we propose that RCOR1 is required for normal myoblast proliferation by inhibiting the expression of p21.

### RCOR1 binds with LSD1 and MyoD in myoblasts and regulates LSD1 stability

RCOR1 is an important binding partner of LSD1 within the CoREST complex and is required for LSD1 catalytic activity and nucleosomal substrate binding^9,18,34^. Therefore, we evaluated LSD1 expression following RCOR1 knockdown and detected a remarkable decrease of LSD1 protein abundance (Fig. 6A), with no effect on LSD1 mRNA expression (Fig. S8A). This was further confirmed by immunofluorescence analysis showing the loss of LSD1 staining signal in the myoblasts following RCOR1 knockdown (Fig. S8B). Upon LSD1 knockout, RCOR1 protein significantly decreased (Fig. 6A), suggesting that RCOR1 and LSD1 interaction is required for the stability of both binding partners. Interestingly, RCOR3 depletion had no effect on LSD1 protein levels (Fig 6A).

**Figure 6.**
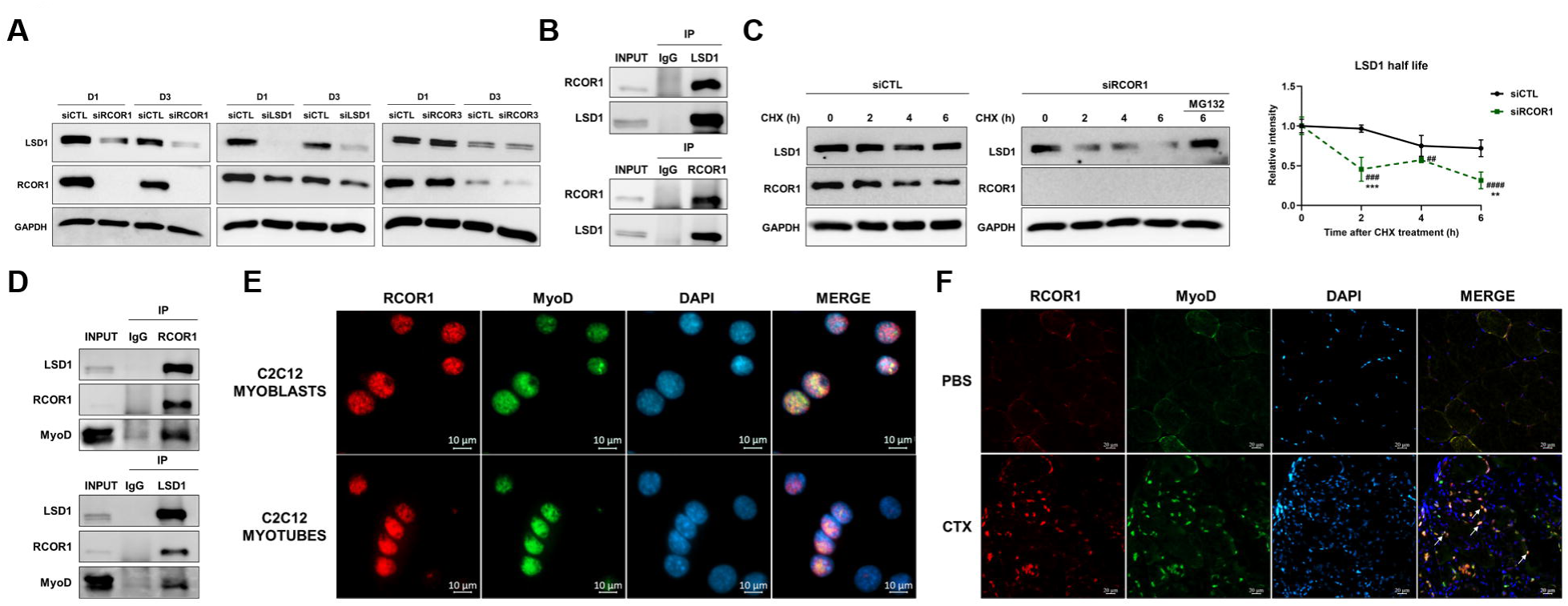
RCOR1 binds with LSD1 and MyoD in myoblasts and promotes/regulates LSD1 stability. C2C12 cells transfected with control siRNA (siCTL), RCOR1, LSD1 or RCOR3 siRNA (siRCOR1; siLSD1; siRCOR3) were cultured in growth medium (GM) and induced to differentiate by switching to differentiation medium (DM) for 3 days. (**A**) Immunoblot was performed to detect protein levels of RCOR1 and LSD1 at day 1 and day 3 in DM. **(B)** C2C12 cells were cultured in GM and induced to differentiate by switching to DM for 1 day. Cell lysates were immunoprecipitated with anti-LSD1 antibody, anti-RCOR1 antibody or rabbit normal IgG. For immunoblotting were used anti-LSD1 and anti-RCOR1 antibodies. (**C**) C2C12 cells were transfected with siCTL or siRCOR1 and 48 hours later, cells were treated with cycloheximide (CHX) (100 µg/mL) for indicated time intervals. siRCOR1-transfected cells were treated with protease inhibitor MG132 (20 μM) in the presence of CHX as indicated for 6 hours before harvest. Immunoblotting was used to detect protein stability and change in the level of LSD1. (**D**) C2C12 cells were cultured in GM and induced to differentiate by switching to DM for 1 day. Cell lysates were immunoprecipitated with anti-LSD1 antibody, anti-RCOR1 antibody or rabbit normal IgG. For immunoblotting was used anti-MyoD antibody. (**E**) C2C12 cells were cultured in GM and induced to differentiate by switching to DM for 3 days. Immunofluorescence staining for RCOR1 (red) and MyoD (green) was performed in myoblasts and myotubes (blue, DAPI). (**F**) Upon cryosection, immunofluorescence staining for RCOR1 (red), MyoD (green) and DAPI (blue) in TA muscles was performed at day 4 following CTX injury. Pictures are one representative of 3 independent experiments. Data are presented as means ± SD, **, P < 0.01; ***, P < 0.001 relative to corresponding siCTL; ###P < 0.01, ####P < 0.0001 relative to siRCOR1 starting point.

To explore whether RCOR1 binds with LSD1 in C2C12 cells, we performed co-immunoprecipitation (Co-IP) experiments. As shown in Fig. 6B, we confirmed the interaction between endogenous RCOR1 and LSD1 indicating that the LSD1-RCOR1 complex is present in C2C12 myoblasts during early stages of differentiation. In order to examine whether RCOR1 affects LSD1 stability we transfected C2C12 myoblasts with siRCOR1 and siCTL and after 48 hours treated the cells with 100 µM CHX to inhibit protein synthesis. Following CHX treatment, endogenous LSD1 drastically decreased over time in siRCOR1 transfected myoblasts relative to control (Fig. 6C). These data indicate that LSD1 protein degraded faster in the absence of RCOR1. Since previous studies suggested that LSD1 is subjected to proteasome degradation^9,35^, siRCOR1 and siCTL-transfected myoblasts were simultaneously treated with CHX and protease inhibitor MG132. Endogenous LSD1 levels remained unchanged following MG132 treatment RCOR1-deficient cells (Fig. 6C).

Upon the initiation of differentiation, MyoD activates the expression of muscle specific genes and the formation of differentiated myotubes. Furthermore, MyoD has been shown to interact with many transcription factors to control muscle gene expression and muscle differentiation^36^. Since RCOR1 knockdown suppressed the expression of known MyoD target genes such as myogenin, we investigated a possible interaction between RCOR1 and MyoD in myoblasts. LSD1 was previously shown to interact with MyoD^12^, allowing us to hypothesize that RCOR1 also associates with MyoD and consequently contributes to the regulation of MyoD-dependent transcription. We performed co-IP experiments during the onset of differentiation in C2C12 cells and confirmed the association between endogenous RCOR1 and MyoD (Fig. 6D). Furthermore, RCOR1 and MyoD were colocalized in the nuclei of C2C12 myoblasts and myotubes (Fig. 6E). MyoD has been reported to be expressed in activated and proliferating SCs upon muscle injury^37^. To assess whether RCOR1 and MyoD are co-expressed in injured muscle, we performed immunofluorescence on regenerating muscle sections. Indeed, the majority of MyoD positive mononucleated cells were also RCOR1 positive, indicating that SCs express RCOR1 as they become activated upon injury (Fig. 6F). Together, these results suggest that MyoD can associate with RCOR1 and LSD1 in C2C12 myoblasts. Both are components of the CoREST complex and this complex could play a role in skeletal muscle differentiation. Furthermore, when RCOR1 is absent, LSD1 becomes more susceptible to proteasomal degradation, suggesting that RCOR1 is required for LSD1 stability in myoblasts.

### Loss of RCOR1 impairs muscle regeneration *in vivo*

To further examine the role of RCOR1 in muscle regeneration *in vivo*, we depleted RCOR1 in the TA muscle during CTX-mediated muscle regeneration. To ensure that RCOR1 was inhibited throughout the experiment, siRCOR1 and siCTL were injected every 2 days into TA muscle during regeneration and muscles were harvested at 10 days for analysis (Fig. 7A). Effective knockdown of RCOR1 protein was confirmed by immunoblot and accompanied with reduction of LSD1 (Fig. 7B), consistent with observations *in vitro* (Fig. 3C and S3D). Both, siRCOR1 and siCTL treated TA muscles displayed newly formed muscle fibers characterized by centralized nuclei (Fig. 7C). However, the number of myofibers containing centrally located nuclei was significantly reduced in siRCOR1 muscles at 10 days following CTX injection (Fig. 7D). We also measured cross-sectional area (CSA) of regenerative myofibers and observed a shift toward smaller sized fibers in TA muscles treated with siRCOR1 (Fig. 7E). Muscle regeneration involves activation and proliferation of SCs and their further differentiation to myocytes before maturing to myofibers. To investigate whether impaired muscle regeneration is associated with reduced pool of satellite cells, we analysed the expression of SC marker Pax7 and found that RCOR1 deficiency in TA muscles inhibited expression of Pax7 (Fig. 7B). Furthermore, differentiation capacity of SCs was also impaired, as shown by supressed expression of myogenic markers MyoD and myogenin in siRCOR1 depleted muscles relative to control (Fig. 7B). These data strongly indicate that RCOR1 contributes to muscle regeneration following injury.

**Figure 7.**
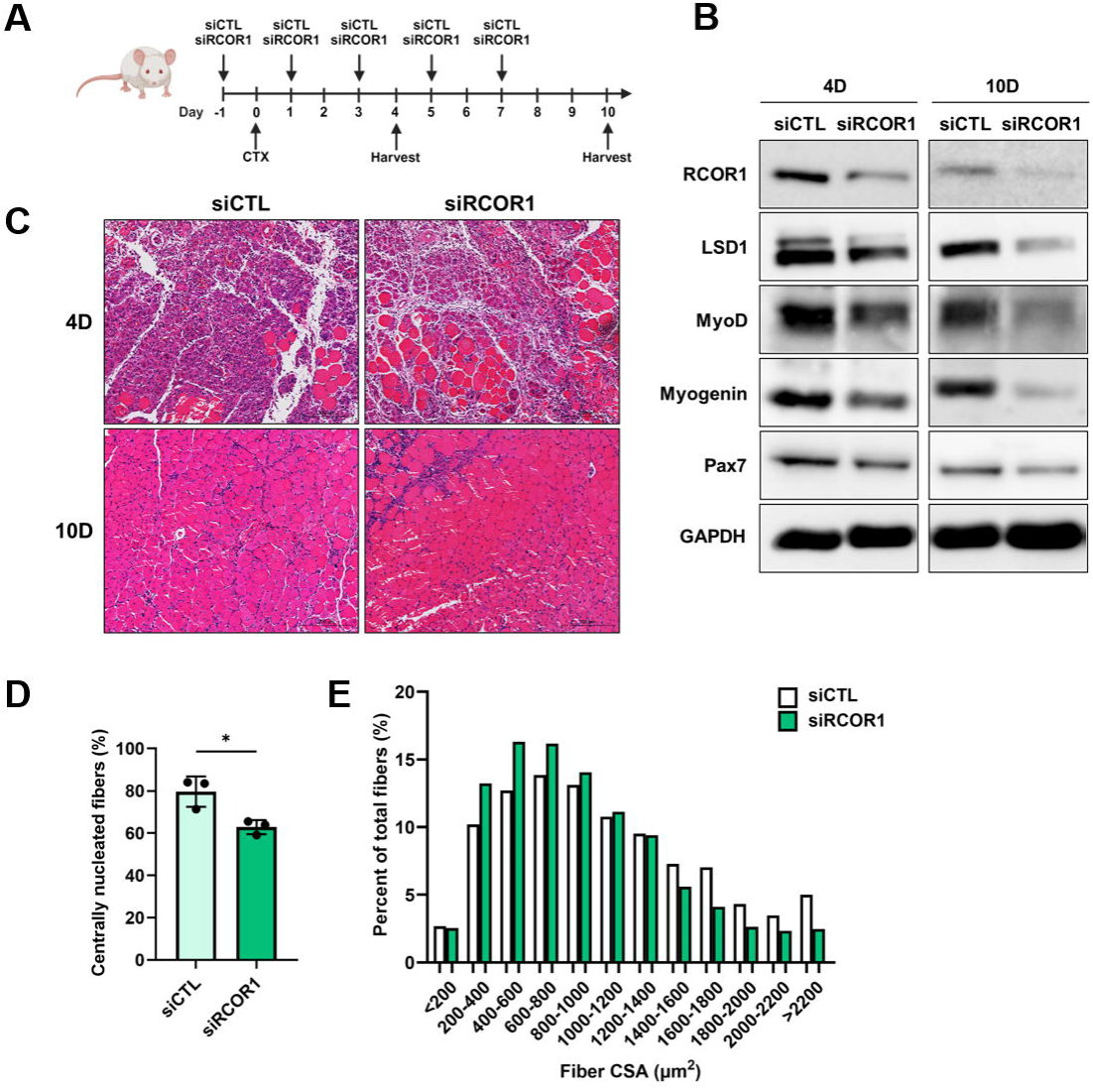
Loss of RCOR1 impairs muscle regeneration. Mouse tibialis anterior (TA) muscles were treated with siRCOR1 or siCTL followed by CTX injection to induce muscle injury. At day 10 following injury, muscles were harvested for analysis. (**A**) Schematic diagram of siRCOR1 mediated knockdown of RCOR1 in the CTX injury muscle model. (**B**) Immunoblot was performed to detect the protein levels of RCOR1, LSD1, Pax7 and myogenic regulatory factors (MRFs) at day 4 and 10 following injury. (C) TA muscles were stained by hematoxylin and eosin (H&E) and analysed for (**D**) percent of centralized myonuclei and (**E**) percent distributions based on CSA (μm2) at day 10. Data are presented as means ± SD; n=3 mice per group. * P < 0.05.

## Discussion

Our study delineates a novel role of RCOR1 as a regulator of myogenesis and muscle regeneration. Loss of RCOR1 remarkably impaired myoblast differentiation *in vitro* and muscle regeneration *in vivo*, caused by reduced number and decreased myogenic aptitude of SCs. Mechanistically, RCOR1 was found to interact with LSD1 and myogenic regulatory factor MyoD, and contribute to LSD1 protein stability in myoblasts. Collectively, our data provide first insights into the role of RCORs, especially RCOR1, in skeletal muscle differentiation.

RCORs are best known as core components of the chromatin-modifying CoREST repressor complex, which also includes LSD1 and HDAC1/2 among several other subunits^23,26,38^. Within this complex, RCOR proteins facilitate LSD1-mediated demethylation of nucleosomes and therefore are essential for LSD1 activity^9,17,34^. During the recent years, LSD1 was identified as a key epigenetic regulator of myoblast differentiation and skeletal muscle regeneration^12,15,16^. This prompted us to investigate whether members of the RCOR family could also have a role in these myogenic mechanisms. RCOR family proteins were initially discovered as transcriptional corepressors for repressor element-1 silencing transcription factor (REST), a key regulator of neurogenesis^23^. Subsequent studies found that RCORs can modulate target gene expression in a REST-independent manner^39^. Among the three members of the RCOR family, RCOR1 exhibits a key role in neurogenesis and regulates hematopoietic differentiation^40,41^. While RCOR2 is involved in the maintenance and regulation of embryonic stem cells (ESCs)^32^ and neurogenesis^40,42^, the role of RCOR3 remains relatively unknown. Although RCORs have been shown to play a role in the development and homeostasis of different tissues, nothing was known about their function in muscle cells. First, we determined that C2C12 cells and primary mouse myoblasts express all three RCOR family members. RCOR1 was expressed at higher level than RCOR2 and RCOR3, which could explain its dominant role in myoblasts. Additionally, RCOR1 formed a complex with LSD1 in myoblasts and protein levels of RCOR1 and LSD1 are found to be interdependent^9,43^. Accordingly, RCOR1 knockdown resulted in a decrease of LSD1 protein in myoblasts, while LSD1 protein levels were unaffected by RCOR3 depletion. These data suggest that the majority of LSD1 protein is found within complexes containing RCOR1 and moreover, RCOR1 was shown to be required for LSD1 stability in myoblasts. Interestingly, LSD1 dysregulation is observed in human patients with neurodegenerative diseases and neuromuscular disorders^44,45^. Thus, targeting LSD1 protein, not only its demethylase activity, could provide some alternative strategies to better target these diseases. Although we have been unable to confirm that RCOR2 or RCOR3 form a complex with LSD1 due to the non-specificity and lack of commercial antibodies, we cannot exclude the possibility that they bind LSD1 in myoblasts.

Previous studies reported that LSD1 levels increase during C2C12 myoblast differentiation^12,15^. LSD1 either physically interacts with MyoD and binds to the myogenin promoter which leads to the activation of myogenic genes or modulates MyoD expression which regulates the entry of myoblasts into the differentiation process^12,15^. Contrary to these reports, we found that both LSD1 and RCOR1 protein levels were highest in proliferating myoblasts and decreased significantly during differentiation to myotubes. As previously mentioned, myogenesis begins with the specification of precursor cells to become myoblasts which further differentiate and fuse to myotubes that ultimately form mature muscle fibers. Tosic et al. showed that LSD1 ablation not only delays skeletal muscle regeneration but also switches the fate of SCs towards brown adipocytes^16^. Thus, it is tempting to suggest that elevated RCOR1 and LSD1 levels in undifferentiated myoblasts are required to restrict the adipogenic potential and direct precursor cells toward the myoblast phenotype. RCOR1 is important for balancing the proliferation and differentiation during brain development, as deletion of RCOR1/2 generated large number of neural progenitors at the expense of differentiated neurons^40^. In addition, a significant decrease of RCOR1 and LSD1 was observed during neuronal differentiation, suggesting that downregulation was necessary for the differentiation process^46^. Given their nuclear localization and similar expression pattern during myoblast differentiation, our data suggest that RCOR1 and LSD1 could act together with MyoD at early stages of differentiation. Indeed, it has been shown that LSD1 associates with MyoD and consequently upregulates myogenin and other muscle-specific genes at the onset of myoblast differentiation^12^. Here, RCOR1 depletion mimicked the effect of LSD1 depletion in myoblasts, as shown by severely impaired myogenic differentiation and downregulated expression of important MyoD target genes such as myogenin, while MyoD remained unaltered. Therefore, we speculated that RCOR1 and LSD1 work together in myogenesis, at least partially through regulating MyoD-dependent transcription activity. Indeed, we confirmed that MyoD binds both RCOR1 and LSD1 in differentiated C2C12 myoblasts, suggesting that they are recruited to the myogenin promoter by MyoD to facilitate muscle-specific gene expression. Most likely reduced LSD1 stability is responsible for the phenotype observed in siRCOR1 myoblasts, suggesting that RCOR1 impacts myoblast differentiation indirectly by altering LSD1 protein level. However, these studies need to be further extended by ChIP experiments to investigate RCOR1 requirement in transcriptional activity of MyoD and whether RCOR1 exerts its function primarily via LSD1.

In addition to the myoblast differentiation defects, RCOR1 knockdown resulted in decreased proliferation and deregulation of many cell cycle genes, without inducing apoptosis. Although myoblast proliferation and differentiation are mutually exclusive^47^, we found both processes to be impaired by RCOR1 depletion, suggesting that they could be regulated through different mechanisms. Consistent with the cell cycle arrest phenotype, RCOR1 knockdown caused an increase of CDK inhibitor p21. Although previous studies showed that p21 is transcriptionally upregulated by LSD1^48^, LSD1 knockdown had no effect on the proliferation rate or the expression of p21 in myoblasts. This discrepancy in proliferation suggests that the role of RCOR1 in cell proliferation might be LSD1-independent and regulated by other subunits of the CoREST complex. Although RCORs have not yet been identified in complexes without LSD1 in mammals, in *Drosophila* the RCOR homolog was present in LSD1 free complexes^49^. Furthermore, LSD1 has also been found in the NuRD^50^ complex, which does not contain RCORs. This complex possesses both histone deacetylase and demethylase activity and coexistence of LSD1/NuRD and LSD1/CoREST complexes has been confirmed. Contrary to RCOR1, RCOR3 depletion resulted in reduced levels of the negative cell cycle regulator p21. RCOR3-deficient myoblasts showed an impaired ability to exit the cell cycle and continued to proliferate under conditions that normally induce differentiation. Although RCOR3 has been previously identified as a natural inhibitor of LSD1^21^, its deletion also inhibited myoblast differentiation by supressing MyoD and myogenin expression, suggesting that RCOR3 might be involved in the transcriptional regulation of MyoD. Indeed, it has been previously proposed that RCORs can target unique genes during cortical development^42^. Equally important was the observation that siRNA-mediated inhibition of RCOR1, RCOR3 and LSD1 resulted in high expression of Cyclin D1, a cell cycle protein whose downregulation is required for the initiation of differentiation^51,52^. Furthermore, inhibition of MyoD function mediated by Cyclin D1 was shown to be unrelated to its role in cell cycle progression^53^.

Notably, RCOR1 depletion in muscle also impaired injury-induced muscle regeneration, as evidenced by decreased size of regenerated myofibers. Muscle regeneration is a highly orchestrated process mediated by the activation, proliferation and differentiation of SCs^54^. Therefore, any dysregulation of these different myogenic stages could lead to impaired regeneration. Here, we found RCOR1 to be highly expressed in MyoD activated SCs during early regeneration phase upon muscle injury, indicating that RCOR1 promotes SC function. Accordingly, RCOR1 depleted muscles exhibited reduced expression of MyoD and myogenin indicating at inhibited myogenic commitment of SCs. This was accompanied with reduced expression of Pax7, which might result from reduced proliferation capacity of SCs in response to RCOR1 depletion, as similar impairment in proliferation was observed in *in vitro* myoblast culture. In addition, we found significantly decreased LSD1 levels following RCOR1 depletion upon muscle injury *in vivo*. Therefore, we propose that both RCOR1 and LSD1 participate in SCs myogenic lineage progression. Interestingly, mice with selective LSD1 ablation in Pax7-positive SCs had no differences in morphology and fiber size distribution, but experienced decreased regenerative capacity upon muscle injury^16^. However, our results demonstrate that LSD1 has an important role in muscle development and postnatal muscle growth, as conditional deletion of LSD1 in limb bud cells resulted in decreased body weight and marked reduction in myofiber size. These phenotype discrepancies are probably due to differences between the analysed mouse models, where LSD1 target genes appear different between skeletal muscle cells and SCs. Previous studies showed that disruption of the RCOR1 gene caused embryonic lethality^40,41^. However, a study with zebrafish RCOR1 mutants showed that these animals managed to survive to adulthood^55^. Although surviving adults developed normally and were fertile, RCOR1 mutants showed locomotive impairment and were hypoactive relative to control animals. Muscle development and regeneration were not evaluated, but these data suggest that RCOR1 knockdown could lead to defects in skeletal muscle in zebrafish. We also analysed mice with germline deletion of RCOR3 (*RCOR3^-/-^*) which seemed healthy with normal progression of skeletal muscle development. However, it is possible that upon injury these mice could display some defects. Given the high sequence similarity of RCOR proteins and their binding ability to LSD1, it is possible that the lack of muscle phenotype in *RCOR3^-/-^* mice is due to a complementary function of the other RCOR family members during muscle development. To fully address this hypothesis, future studies with a conditional double knockout of RCOR1 and RCOR3 would be required to circumvent the potential for redundancy. Also, further analysis will be needed to understand the relevance of RCOR3 in skeletal muscle regeneration. Importantly, RCOR2 and RCOR3 mRNA expression upregulated following muscle injury, but remained at very low and almost undetectable levels. Taken together, these data suggest that RCOR1 is required for activation and differentiation of SCs during muscle regeneration.

These data are the first evidence for a unique role of RCOR1 in skeletal muscle cells and muscle regeneration. RCOR1 was shown to function in myoblast proliferation, differentiation and fusion into myotubes by regulating the expression of cell cycle and muscle-specific genes. Moreover, RCOR1 depletion decreased injury-induced SC activation and differentiation resulting in impaired muscle regeneration. Although, the molecular mechanisms by which RCOR1 regulates myogenesis are not fully characterized and remain to be further studied, our findings provide new insights into potentially exploitable therapeutic factors in musculoskeletal health.

## Materials and Methods

### Animals

Mouse studies were approved by the Finnish ethical committee for experimental animals (license number #14044/2020), complying with the international guidelines on the care and use of laboratory animals. Mice on a C57BL/6JRj background were housed under standard laboratory conditions and were fed ad libitum. All mice used for analyses were male. The mice were euthanized with CO2-asphyxiation followed by sample collection. For the analysis of RCOR1 expression during postnatal muscle development, hindlimb muscles were obtained from mice at postnatal day 1 (P1), 2 weeks (W2) and 4-6 weeks (W4-6) of age. Neonate mouse pups were euthanized by decapitation. Mice embryos carrying the floxed Kdm1a/Lsd1 alleles (Lsd1^fl/fl^, stock #023969) were obtained from the Jackson laboratory. Lsd1^fl/fl^ mice were crossed with Prrx1-Cre to create limb bud mesenchyme specific conditional Lsd1-knockout mice (LSD1^Prrx1-/-^), as described previously^31^. Mice were analysed at 4 weeks of age.

### Generation of mice with a germline deletion of RCOR3^-/-^

The murine embryonic stem (ES) cells carrying targeting vector for the RCOR3 gene were obtained from the European Mouse Mutant Cell Repository (EuMMCR). The ES cells were injected into C57BL/6N mouse blastocysts to generate chimeric mice. Heterozygous Rcor3^LacZ/+^ mice were bred with CAG-Cre mice expressing Cre recombinase under the control of CMV-IE enhancer/chicken β-actin/rabbit β-globin hybrid promoter in widespread tissues, leading to deletion of exons 4 and 5 of the RCOR3 gene and generation of *RCOR3^+/-^* mice (Fig. S6A and B). *RCOR3^-/-^* animals were obtained from breeding heterozygote mice, and WT littermates (*RCOR3^+/+^*) were used as controls. Genotyping of mice was carried out from DNA extracted from ear marks of 2-week-old to 3-week-old mice. Mice were analysed at 6 weeks of age. The primer sequences used for the PCRs are shown in Supplementary Table S1.

### Cell culture

For all experiments, C2C12 myoblasts (ATCC) were cultured in high-glucose Dulbecco’s Modified Eagle Medium (DMEM; Sigma-Aldrich), supplemented with 10% foetal bovine serum (FBS; Gibco) and 1% penicillin/streptomycin (Gibco) (Growth Medium, GM). Once cells reached 80-90% confluence, differentiation was induced by DMEM supplemented with 2% horse serum (HS, Gibco) and 1% penicillin/streptomycin (Differentiation Medium, DM). The medium was changed every other day.

Primary myoblasts were isolated from the hindlimbs of 8±1 week-old C57BL/6JRj male mice as described previously^56^. Briefly, muscles were minced using scissors and digested with 400 U/ml of collagenase II (Gibco) solution for 1 h at 37 °C. To remove tissue debris and large infiltrating cells, the digested tissue suspension was passed through a 70 µm cell strainer and an additional 30 µm filter. Cells were plated on 10 cm Matrigel coated dishes in Ham’s F10 nutrient mixture (Sigma Aldrich) with 20 % FBS, 10 ng/mL fibroblast growth factor 2 (FGF2; PeproTech) and 1% penicillin-streptomycin. To remove fibroblasts, cells were incubated with a small amount of PBS at room temperature (RT) for 5 min. The detached cells were collected with Ham’s F10 nutrient mixture and plated on a new Matrigel coated dish. This was repeated until no fibroblasts were observed in the culture. Differentiation was induced with DMEM supplemented with 5% HS and 1% penicillin-streptomycin when cells reached 90% confluency. The medium was changed every other day. All cells were cultured in a 37°C incubator with 5% CO2.

### siRNA-mediated gene knockdown

For knockdown experiments, C2C12 cells and primary mouse myoblasts were transfected with SMARTpool siRNA oligonucleotides (Dharmacon) using Lipofectamine RNAImax (Invitrogen), according to the manufacturer’s instructions. Briefly, RNAImax-siRNA complexes were added into each well to yield a final concentration of 20 nM siRNAs. Differentiation was induced 24 hrs after and cells were harvested at indicated time points. The siRNAs used in this study were SMARTpools of ON-TARGETplus Non-targeting siRNA (siCTL), ON-TARGETplus Mouse LSD1 siRNA (siLSD1), ON-TARGETplus Mouse RCOR1 siRNA (siRCOR1), ON-TARGETplus Mouse RCOR2 siRNA (siRCOR2) and ON-TARGETplus Mouse RCOR3 siRNA (siRCOR3) (Dharmacon). The efficiency of mRNA and protein knockdown was confirmed using qPCR and immunoblotting, respectively.

### Cell proliferation assay

Cell proliferation was measured by CellTiter 96® Non-Radioactive Cell Proliferation Assay kit (MTT; Promega) according to the manufacturer’s instructions. Briefly, C2C12 myoblasts were plated at 2000 cells per well in 96-well plates and assayed at 24, 48- and 72-hours post plating. At indicated times, 15 µl of dye solution was added directly to the wells and plates were incubated for 3 hours in a humidified 37 °C incubator with 5% CO2. Absorbance was measured at 570 nm by using an absorbance microplate reader (Ensight).

### 5-Bromo-2-deoxyuridine (BrdU) assay

5-Bromo-2-deoxyuridine (BrdU) assay was performed with BrdU (Santa Cruz Biotechnology) and anti-BrdU antibody (Santa Cruz Biotechnology). Cells transfected with siCTL, siLSD1, siRCOR1 and siRCOR3 were cultured in growth medium for 24 hours and then switched into fresh growth medium or differentiation medium. After 24 hours cells were stimulated with growth or differentiation medium containing 10 µM BrdU for 4 hours prior to harvesting. Cells were fixed, permeabilized, treated with 1M HCl for 1 hour and incubated overnight at 4°C with anti-BrdU antibody following incubation with Alexa Fluor-488 conjugated goat anti-mouse IgG (Jackson ImmunoResearch Laboratories). Slides were mounted in mounting medium for fluorescence with DAPI (Vector Laboratories). The proliferation rate was calculated as the ratio between cells stained with BrdU and those stained with DAPI.

### Apoptosis assay

C2C12 myoblasts were stained with Annexin V FITC/PI kit (Santa Cruz Biotechnology) according to the manufacturer’s instructions. Briefly, cells were incubated with 500 μL 1x binding buffer, 2.5 μg FITC-Annexin V and 50 µl propidum iodide (PI) at RT for 15 min in the dark. The stained samples were detected by fluorescence microscopy.

### RNA isolation and RT-qPCR

Total RNA was isolated using NucleoSpin RNA Plus kit (Macherey-Nagel) following the manufacturer’s instructions. cDNA synthesis was performed with Sensifast cDNA synthesis kit (Meridian Bioscience) and Quantitative PCR (qPCR) on CFX384 Real-Time PCR Detection System using DyNAmo Flash SYBR Green qPCR Kit (ThermoFisher Scientific). Relative quantification of gene expression was performed by the 2^−ΔΔCt^ method using Gapdh as the housekeeping gene for normalization. Primer sequences are shown in Table S1.

### Immunoblotting

Cells were lysed in radioimmunoprecipitation assay (RIPA) buffer (50 mM Tris-HCl, pH 7.4; 0.5% NP-40; 0.25% sodium deoxycholate; 150 mM sodium chloride; 1 mM sodium orthovanadate; 0.5 mM phenylmethylsulfonyl fluoride), while muscle tissues were lysed in Lysis buffer (20 mM Tris-HCl, pH 7.8; 137 mM NaCl; 2,7 mM KCl; 1 mM MgCl2; 1% Triton x-100; 10% glycerol; 1 mM EDTA, 1 mM DTT), both supplemented with protease inhibitor cocktail (Roche). Samples were incubated on ice for 30 min and cell debris was separated by centrifugation at 14 000 rpm for 15 min at 4 °C. Supernatant was collected and protein concentration was determined by Bradford colorimetric assay (Bio-Rad). Equal amount of protein was loaded on 12% SDS-PAGE gels and transferred to nitrocellulose membranes. Blocking was performed in Tris-buffered saline with 5% non-fat dry milk for 1 h at RT. Membranes were probed with primary antibodies at 4°C overnight, followed by incubation with horseradish peroxidase (HRP)-linked secondary antibodies (Cell Signaling) for 1 hour at RT. The blots were visualized with WesternBright Quantum kit (Advansta) and images were taken with the LAS-4000 Luminescent imager (Fujifilm Life Sciences). Quantification of the density of each band was performed by Image J software. Antibody information can be found in Table S2.

### Immunofluorescence

At indicated time points, cells were rinsed in PBS, fixed in 4% formaldehyde (PFA, Invitrogen) for 15 min and permeabilized in 1% TritonX-100/PBS solution for 20 min at RT. Cells were then blocked for 30 min in PBS/ 3%BSA/ 0.05%Triton X-100 blocking solution and incubated with primary antibodies prepared in blocking solution for 2 h RT or overnight at 4 °C. This was followed by 3 washes in PBS and incubation with Alexa Fluor-594-conjugated goat anti-rabbit IgG and Alexa Fluor-488 conjugated goat anti-mouse IgG (Jackson ImmunoResearch Laboratories) for 1h in the dark. After washing in PBS, slides were mounted in mounting medium for fluorescence with DAPI (Vector Laboratories). Images were obtained by Axio Imager Z2 (Zeiss) microscope using ZEN software.

### Co-immunoprecipitation experiments

Cells were washed in PBS, lysed in IP buffer (10 mM Tris pH 8, 0.4% NP40, 300 mM NaCl, 10% glycerol, 1 mM DTT, protease inhibitor cocktail (Roche)), passed 10 times through a 25G needle and incubated for 30 min at 4 °C. Cell lysates were centrifuged at 15,000 × g for 15 min at 4 °C and the supernatants were diluted with one volume of dilution buffer (10 mM Tris pH 8, 0.4% NP40, 5 mM CaCl2). Samples with equal amounts of proteins were precleared with Dynabeads Protein G (Thermo Fisher Scientific) for 2 hours at 4 °C and then incubated with 3 µg of primary antibody on a rotary platform overnight at 4 °C. The next day, beads were blocked in 5 mg BSA/PBS for 2 h at 4 °C and then added to the samples for additional 2 hours at 4 °C. The unbound supernatant was aspirated and the beads were washed with ice-cold diluted IP buffer. Proteins were eluted with SDS sample buffer by heating at 99 °C for 5 min and analysed by immunoblotting.

### Protein stability assay

C2C12 cells were transfected with 20 nM concentration of siCTL and siRCOR1. After 48 hours cells were treated with 100 μg/ml cycloheximide (CHX; Sigma Aldrich) to inhibit protein synthesis. CHX-treated cells were harvested at different time points (0, 2, 4 and 6 hours) and processed for immunoblotting with anti-LSD1 (Invitrogen) and anti-RCOR1 (Abcam) antibody. Anti-ß-actin antibody (Abcam) was used as internal control. Subsequently, C2C12 cells were simultaneously treated with proteasomal inhibitor MG132 (20 μM: Santa Cruz Biotechnology) for 6 hours and processed for immunoblotting.

### Cardiotoxin injury

Cardiotoxin (CTX; Latoxan, France) was dissolved in sterile saline to a final concentration of 100 µM. Male 8-10 weeks old C57BL/6JRj mice were anesthetized with isoflurane inhalation and provided with analgesia (0.1 mg/kg buprenorphine) to alleviate pain. Right tibialis anterior (TA) muscles were intramuscularly injected with 20 μl of 20 µM CTX solution with a hypodermic needle, while left TA muscles were injected with sterile saline only. Regenerating TA muscles were isolated 4, 7, and 10 days following CTX injection.

### Intramuscular transfection of siRNAs

Intramuscular transfection of siRNAs (5 µg) was performed using *in vivo*-jetPEI kit (Polyplus) following the manufacturer’s recommendations. PEI/siRNA complexes were formed at an N/P ratio of 8 in equal amounts of 5% glucose solution. After 15 minutes incubation at RT, PEI/siRNA complex mixture (20 µl) was injected intramuscularly into mouse TA muscles with a hypodermic needle. Hindlimbs of 8-10-week-old male mice were shaved and cleaned with 75% alcohol. The mixture containing si-RCOR1 was injected into the right TA muscle, while the mixture containing si-CTL was injected into the left TA muscle as a negative control. Regenerating TA muscles were isolated 4, and 10 days following CTX injection.

### TA muscle histology

For histological analyses, freshly isolated TA muscles were fixed in formalin, dehydrated by graded ethanol, embedded in paraffin, and cut into 5 μm sections. The sections were deparaffinized, rehydrated, and stained with haematoxylin & eosin. Cross-sectional areas (CSA) and diameters of TA fibers were measured using ImageJ software. Minimum 1,000 muscle fibers were measured for each sample. For immunofluorescence studies, regenerating TA muscles were flash-frozen in liquid nitrogen-cooled isopentane. Frozen blocks were embedded in Optimal Cutting Temperature (O.C.T.) compound and sectioned at 8 μm using a cryostat microtome. Following fixation with 4 % formaldehyde for 15 minutes, antigen retrieval was performed by incubating muscle sections in boiling 10 mM citrate buffer pH 6 for 10 minutes. Then, samples were permeabilized with 0.5 % Triton X-100 for 20 minutes, blocked with 3 % BSA for 1 hour and incubated with AffiniPure Fab Fragment Goat Anti-Mouse IgG (H + L) (Jackson ImmunoResearch) in PBS (20 µg/mL) for 1 hour. Primary antibodies were incubated overnight at 4 °C, followed by incubation with Fluor Alexa 488- and 594-conjugated secondary antibodies (Jackson ImmunoResearch Laboratories). DAPI was used to visualize nuclei. PBS washes were performed between each step. Unless indicated otherwise, all incubations were carried out at RT.

### Quantification and Statistics

The differentiation index was calculated as the percentage of nuclei in MyHC-positive cells compared to the total number of nuclei. The fusion index was calculated as the percentage of nuclei in MyHC-positive myotubes with ≥ 2 nuclei out of the total nuclei. All data were expressed as mean ± standard deviation (SD). Statistical differences between groups were determined by unpaired Student’s t-test and one-way ANOVA with Tukey’s post-hoc test by using GraphPad Prism 10.2.0. All experiments were performed independently at least three times. P values smaller than 0.05 were considered to be significant (*p < 0.05; **p< 0.01; ***p< 0.001; ****p< 0.0001).

## Acknowledgements

The authors thank Merja Lakkisto and the staff of Turku Central Animal Laboratory for their excellent technical assistance.

## Author contributions

MP and RK designed and conceived the study; RK supervised the study; MP, FW, PR and MR performed experiments and analysed data; RK and HT provided advice and technical assistance; and MP wrote the manuscript. All authors have contributed to and approved the final manuscript.

## Data Availability

The manuscript does not contain RNA-Seq or other large data sets. Detailed information on antibodies and primers used in the manuscript is provided in the supplemental tables.

