## Supplementary figures and images for "RCOR1 promotes myoblast differentiation and muscle regeneration"

### Supplementary Figures and Tables

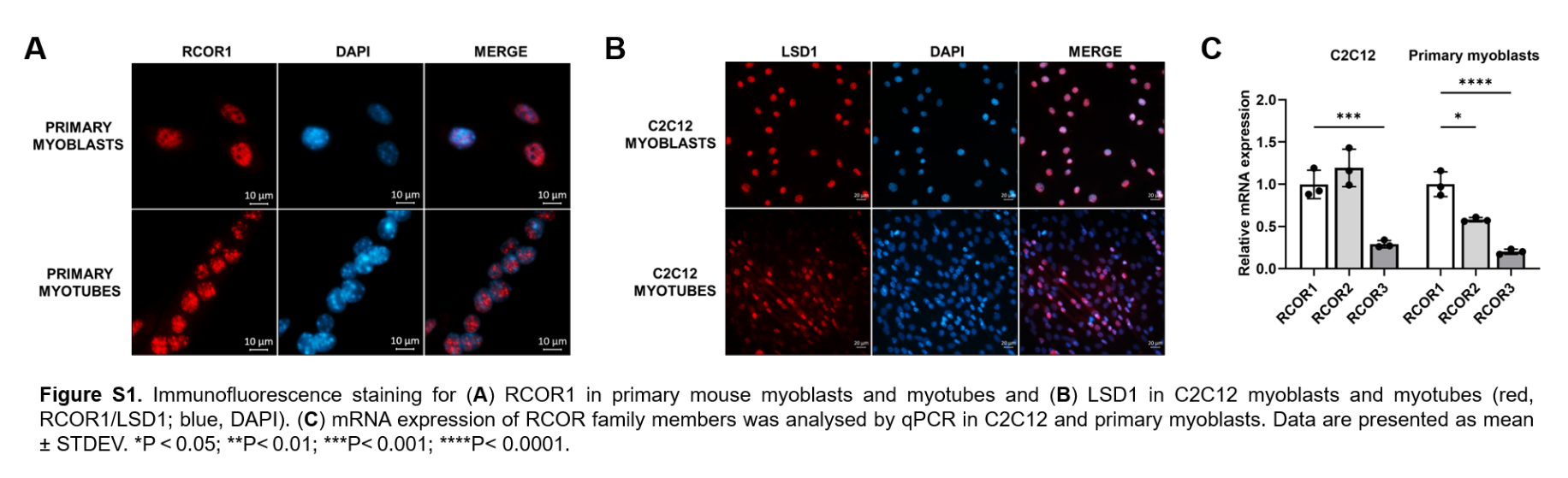

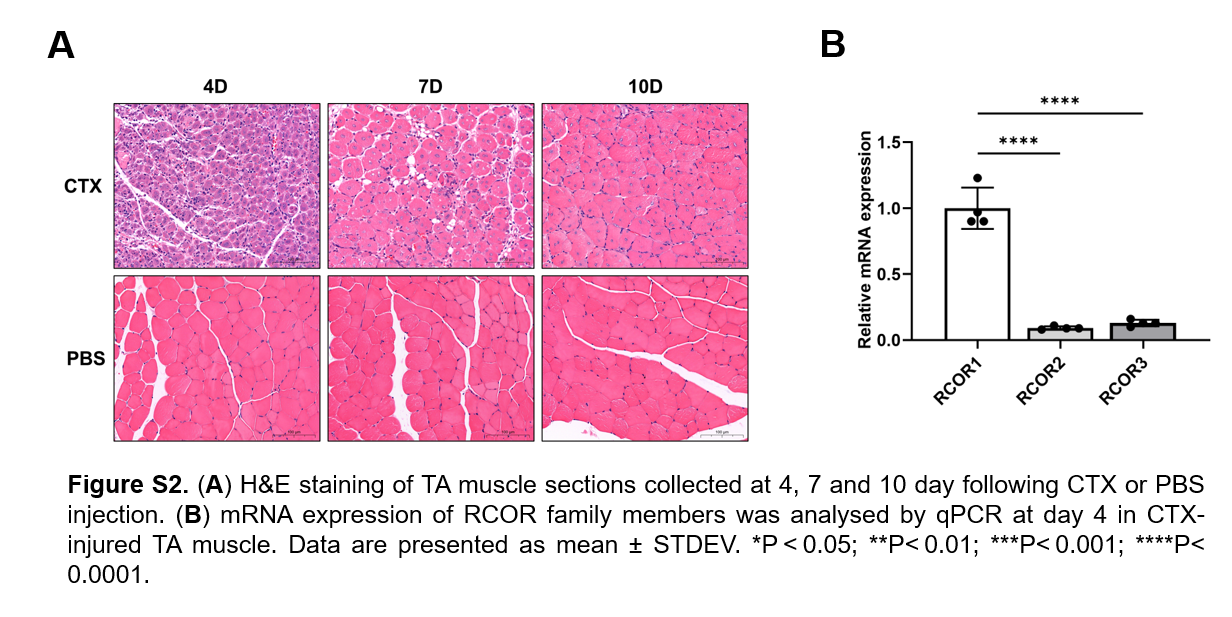

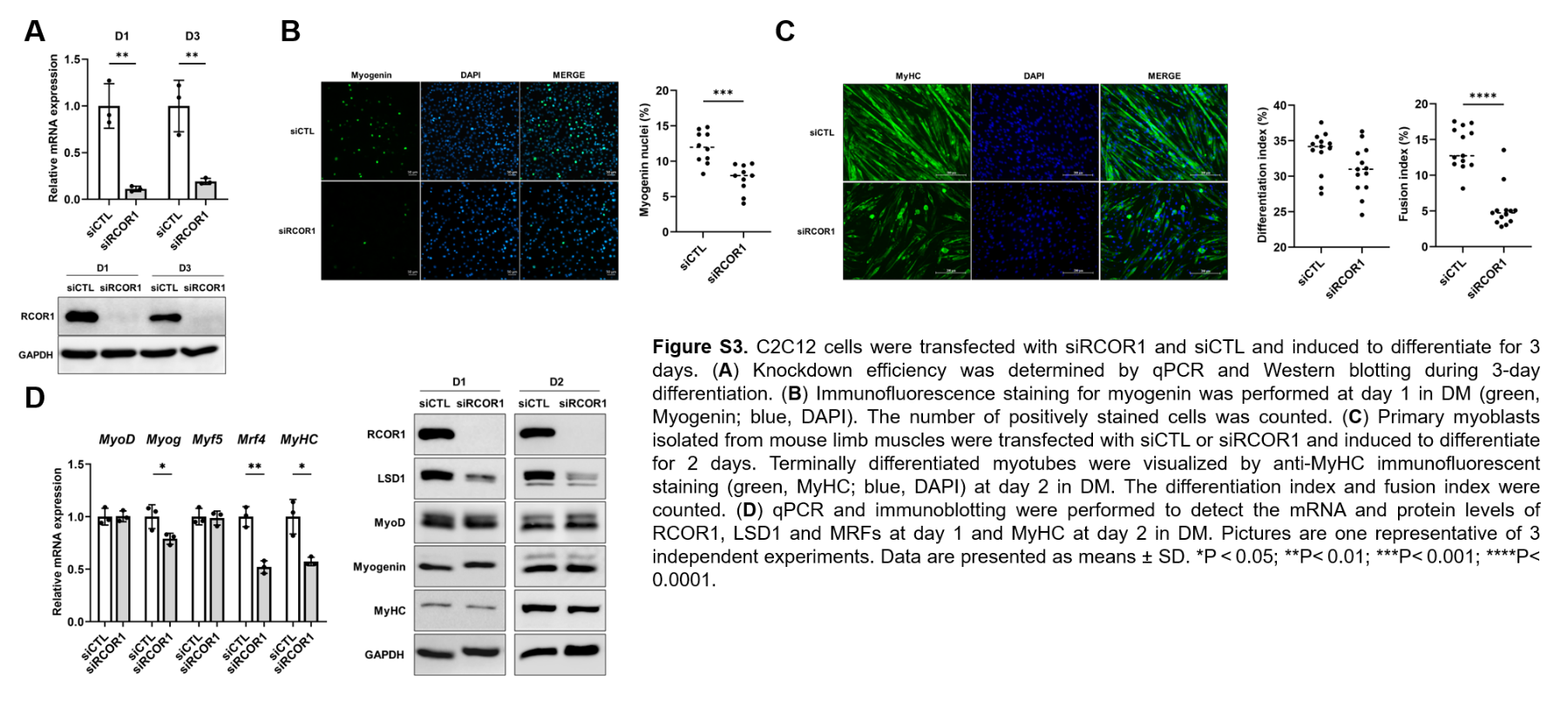

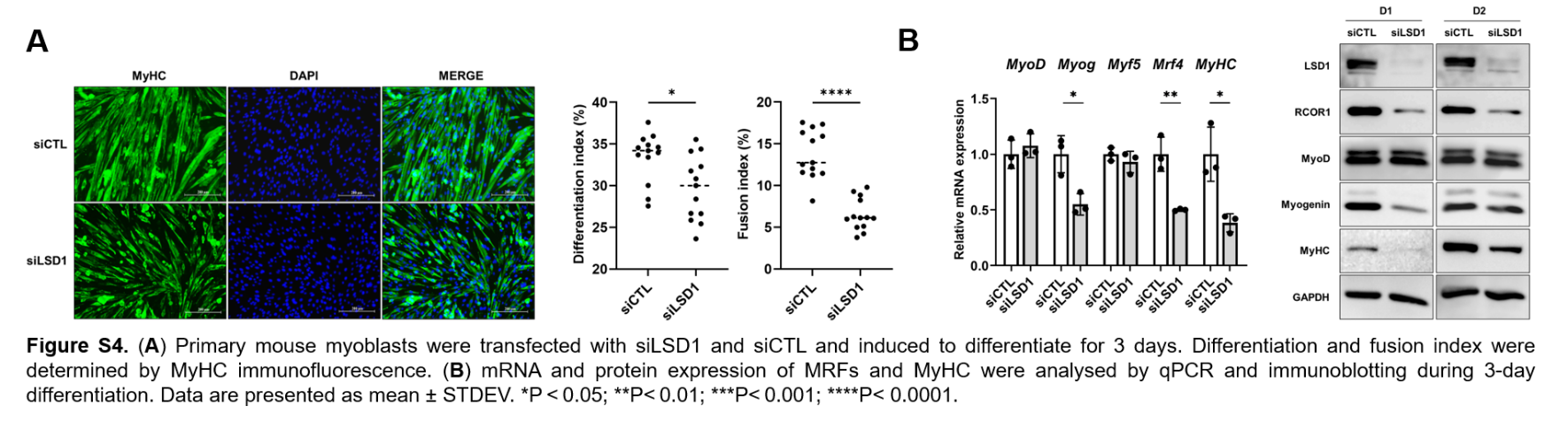

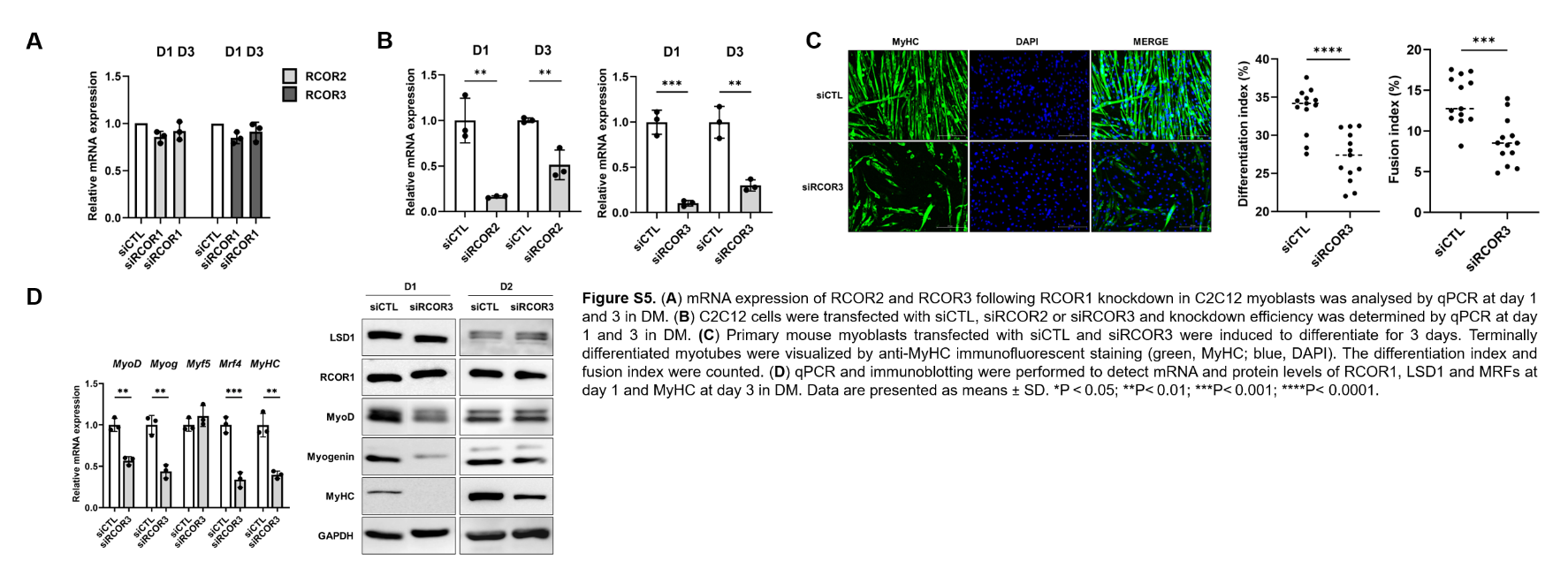

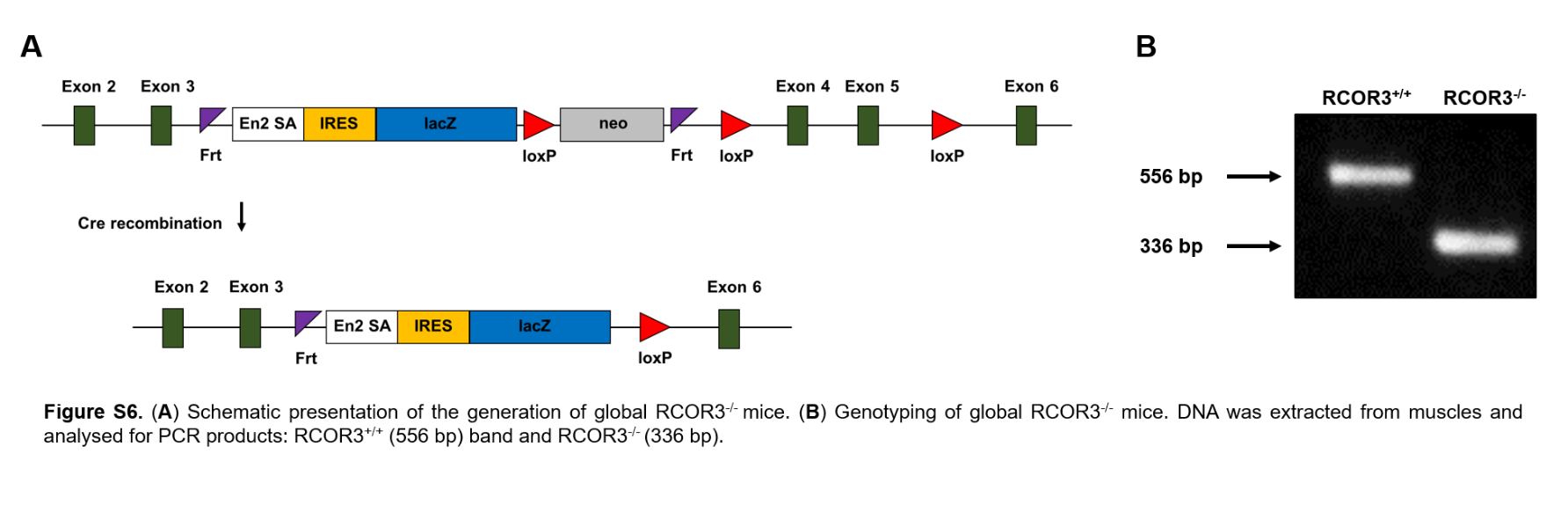

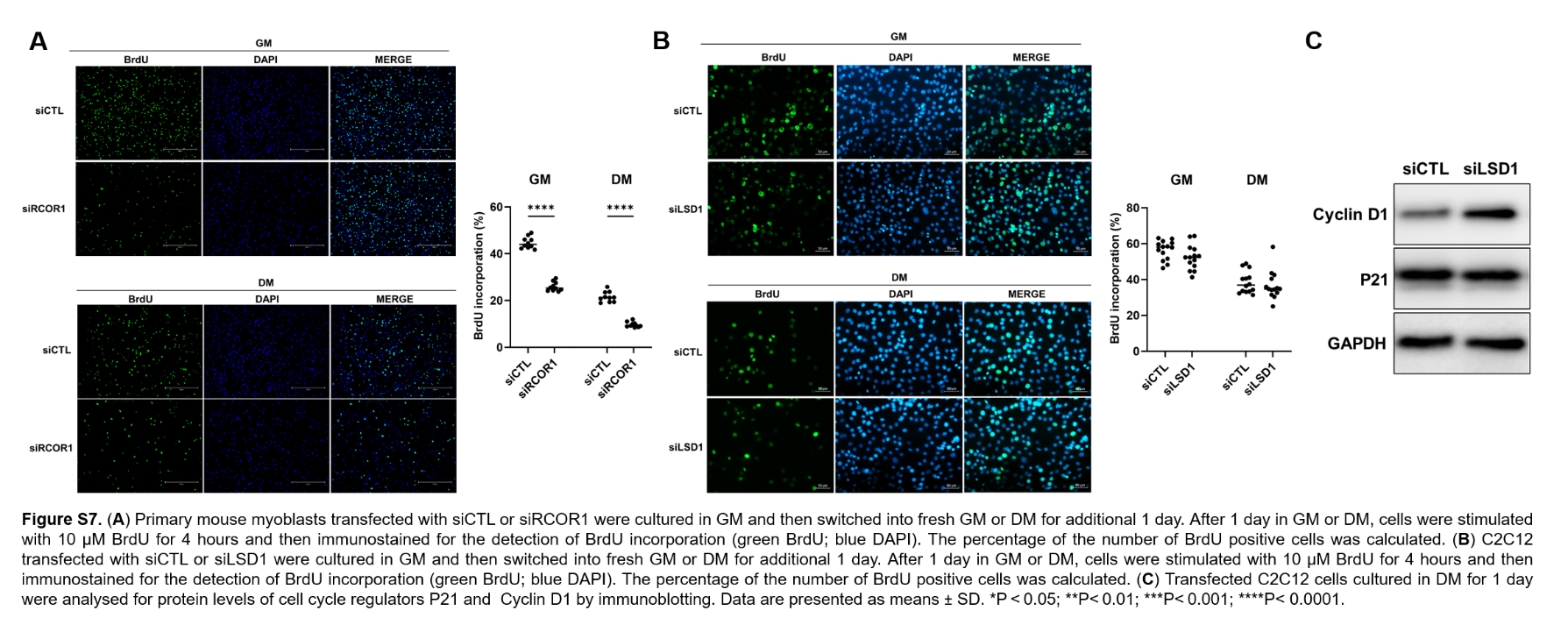

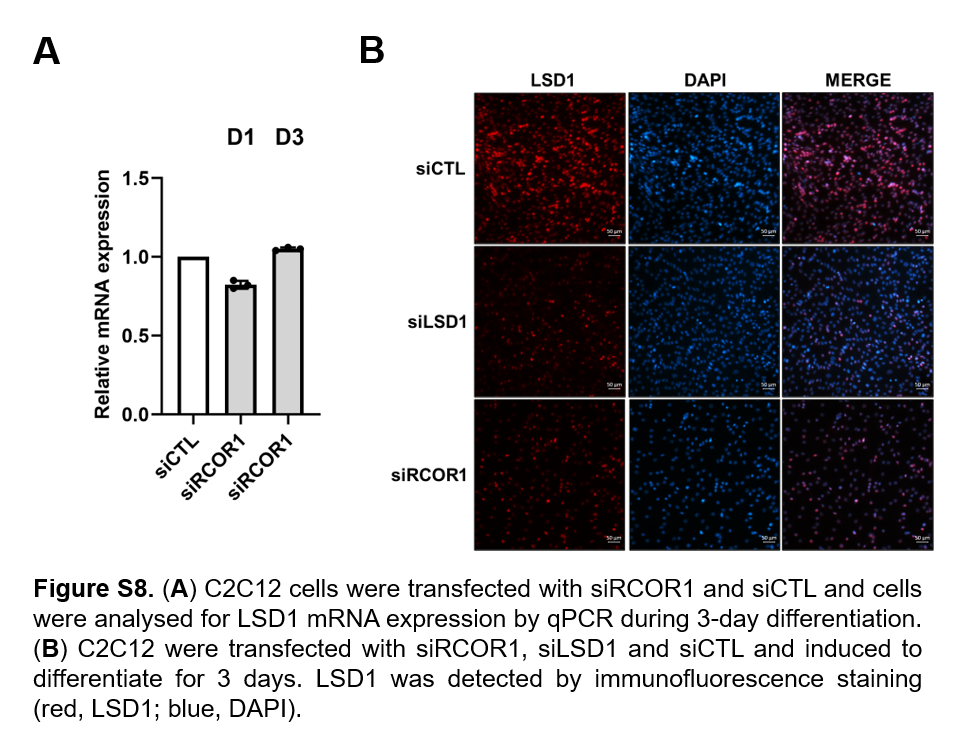

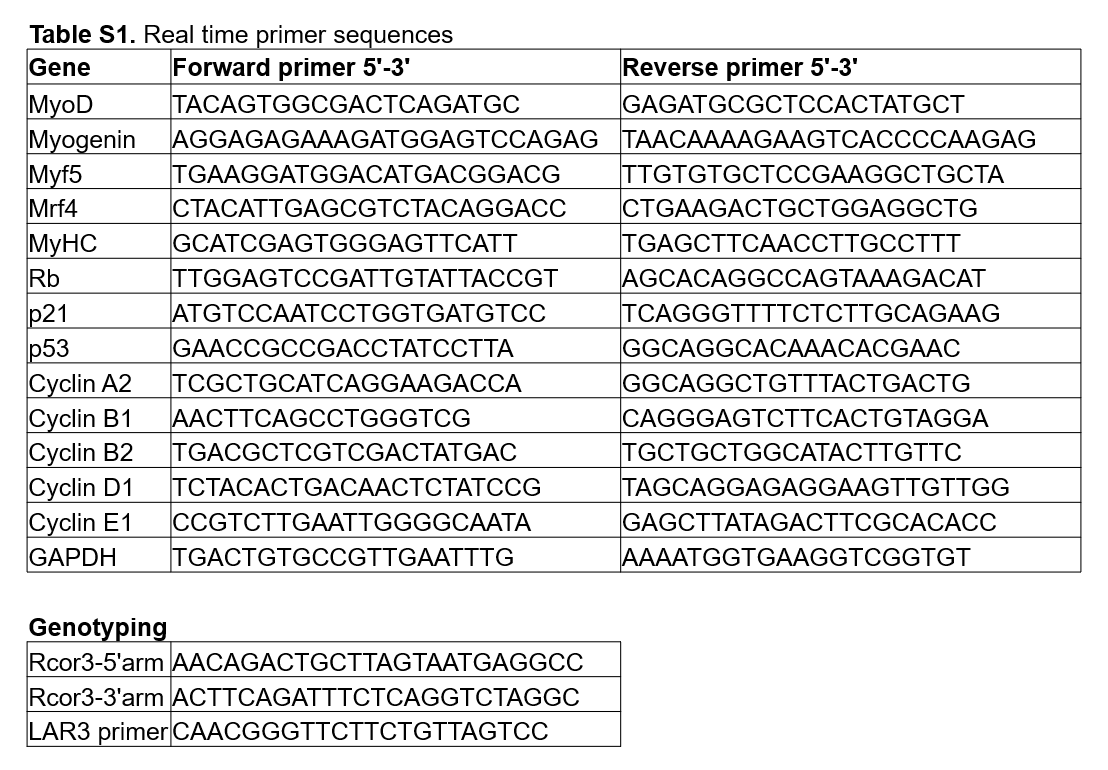

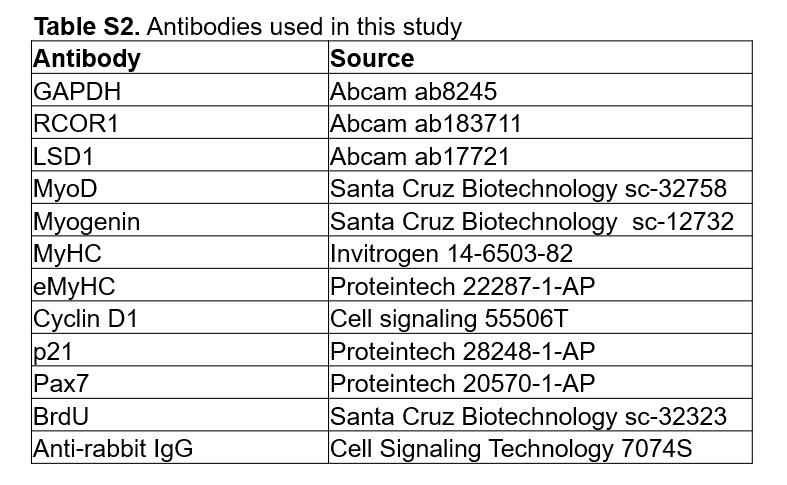
